# Uncovering the Genomic Manifold via Scalable Learning from the Global Microbiome

**DOI:** 10.1101/2025.01.30.635558

**Authors:** Weimin Wu, Zhihan Zhou, Robert Riley, Mohammad Abdulqader, Xuefeng Song, Satria Kautsar, Rob Egan, Steven Hofmeyr, Gefei Liu, Shira Goldhaber-Gordon, Mutian Yu, Harrison Ho, Yaping Liu, Andrei Stecca Steindorff, Fengchen Liu, Feng Chen, Rachael Morgan-Kiss, Lizhen Shi, Han Liu, Zhong Wang

**Affiliations:** Northwestern University, Evanston, IL, USA; Lawrence Berkeley National Laboratory, Berkeley, CA, USA; Brown University, Providence, RI, USA; Johns Hopkins University, Baltimore, MD, USA; University of California at Merced, Merced, CA, USA; University of California at Berkeley, Berkeley, CA, USA; Miami University, Oxford, Ohio, USA; Illumina Inc., Foster City, CA, USA

**Keywords:** GenomeOcean, Genomic Manifold, Metagenomics, Genome Foundation Model

## Abstract

Functional genomic sequences occupy an infinitesimal fraction of the astronomically large DNA sequence space, implying that evolution explores a low-dimensional manifold shaped by universal biochemical and evolutionary constraints. To investigate this underexplored **Genomic Manifold Hypothesis** using scalable, data-driven evidence, we developed **GenomeOcean**, a 4-billion-parameter generative model trained on 219 TB of diverse, co-assembled global environmental metagenomic data using an optimized transformer. The model captures both universal phylogenetic syntax and protein-coding fidelity. Furthermore, we validate the embeddings’ intrinsic low dimensionality. Crucially, comparisons with the independently trained Evo 2 model demonstrate a strong linear correspondence between their embedding spaces and convergent generative behaviors. These results suggest that the genomic manifold represents a fundamental biological principle, offering a unified framework for understanding evolutionary constraints and guiding synthetic biology.

## 1 Introduction

Functional genomic sequences occupy an infinitesimal fraction of the astronomically large DNA sequence space, theoretically represented by 4*^N^* permutations for a sequence of length *N* (Axe, 2004; Karas and Hecht, 2020; Pevsner, 2015). For instance, estimates suggest that fewer than 1 in 10^77^ random sequences can adopt a functional protein fold (Axe, 2004). This extreme sparsity implies that evolution does not explore sequence space randomly, but is constrained to a low-dimensional, structured subspace: the **Genomic Manifold**. Shaped by universal biochemical laws and fitness landscapes, this manifold constitutes a conserved topological structure inherent to all living systems. While the existence of such a manifold is a fundamental biological postulate, scalable and data-driven evidence for its geometry has remained elusive. Traditional analyses, constrained by phylogenetically biased reference genomes, fail to capture the full global diversity required to map this continuous structure (Albright and Louca, 2023; Ding and Steinhardt, 2024).

Foundation models offer a transformative approach to this challenge by learning generalizable structures directly from sequence data (Brixi et al., 2025; Dalla-Torre et al., 2023; Nguyen et al., 2024; Zvyagin et al., 2022). Much like large language models that map the semantic topology of human language (Brown et al., 2020; Dubey et al., 2024; Goldstein et al., 2024; Jiang et al., 2024; Kenton and Toutanova, 2019; Park et al., 2024; Zada et al., 2025), genome foundation models (gFMs) have the potential to uncover the latent syntax of biology. However, current gFMs are largely trained on reference genomes, creating a discontinuous sampling bias that represents the continuous spectrum of evolution as a set of isolated points (Brixi et al., 2025; Dalla-Torre et al., 2023; Ji et al., 2021; Li et al., 2025; Liu et al., 2025; Nguyen et al., 2023, 2024; Sanabria et al., 2024; Schiff et al., 2024; Wu et al., 2025; Zhou et al., 2023; Zvyagin et al., 2022). This discrete sampling is insufficient for learning a manifold, as the model cannot infer the valid evolutionary paths, or “gradients”, that connect these isolated species without observing the intermediate sequences. By relying on curated isolates, these models exclude “microbial dark matter”: the vast, uncultured diversity that constitutes the majority of Earth’s genetic information (Rinke et al., 2013). They also omit the “rare biosphere”, consisting of low-abundance populations that play crucial ecological roles (Pascoal et al., 2021; Sogin et al., 2006). Lacking these information, reference-based models likely perceive the genomic manifold as fragmented rather than continuous. Therefore, to accurately reconstruct the geometry of the genomic manifold, modeling must extend beyond sparse reference databases to capture the full, uncurated continuity of the global microbiome.

Here, we introduce **GenomeOcean**, a 4-billion-parameter generative foundation model trained on 645.4 Gbp of high-quality contigs assembled from over 219 TB of raw metagenomic reads. GenomeOcean distills this vast environmental information through large-scale co-assembly (Har, 2018; Oliver et al., 2024; Peterson et al., 2009; Riley et al., 2023; Sunagawa et al., 2020; Wang et al., 2019), recovering the structural connectivity of the genomic manifold that reference databases omit. Leveraging byte-pair encoding (BPE) and an optimized transformer decoder architecture (Ainslie et al., 2023; Dao, 2023; Elfwing et al., 2018; Jiang et al., 2024; Sennrich et al., 2016; Su et al., 2024; Vaswani et al., 2017; Zhang and Sennrich, 2019), the model reveals a comprehensive view of latent genomic structure.

Our analysis demonstrates that GenomeOcean simultaneously encodes coarse-grained universal syntax (phylogenetic relationships) and fine-grained evolutionary fidelity (protein-coding constraints). The model generates full-length protein-coding regions that fold into native-like structures, even when trained primarily on fragmented metagenomic contigs. Furthermore, we validate that the high-dimensional GenomeOcean embeddings possess an intrinsic low-dimensional structure. Crucially, comparisons with the independently trained Evo 2 model (Brixi et al., 2025) reveal a strong linear correspondence between their embedding spaces and convergent generative behaviors. This convergence suggests that the genomic manifold is not a model artifact, but a robust biological reality. Together, these findings provide large-scale, data-driven evidence that Earth’s genetic diversity inhabits a shared, navigable design space, offering a unified framework for understanding evolutionary constraints and guiding synthetic biology, such as mining novel biocatalysts, engineering enzymes from uncultured organisms, or designing metabolic pathways.

## 2 Results

### A Global Metagenomic Corpus for Modeling

To probe the geometry of the genomic manifold while minimizing the potential biases of reference genomes, we first compiled a global sampling of environmental metagenomes (**Figure** 1**A**). Current genome foundation models are often trained on reference genomes, which may fail to capture the vast, uncultured diversity of the rare biosphere. It is estimated that the Earth contains an estimated one trillion species of microbes — with 99.999% of them remaining undiscovered (Locey and Lennon, 2016). To overcome this limitation, we applied a large-scale co-assembly strategy to several environmental metagenomics datasets (**Methods** 4.2.1). As shown previously, the co-assembly strategy effectively recovers species that are missed in single-sample assemblies (Hofmeyr et al., 2020; Riley et al., 2023). Using the resulting contigs, instead of metagenome-assembled-genomes (MAGs), which, in part, rely on curated reference databases of conserved gene markers for quality assessment (Parks et al., 2015, 2022), this strategy increases the representation of low-abundance species, which are often omitted during the metagenome binning process.

**Figure 1.**
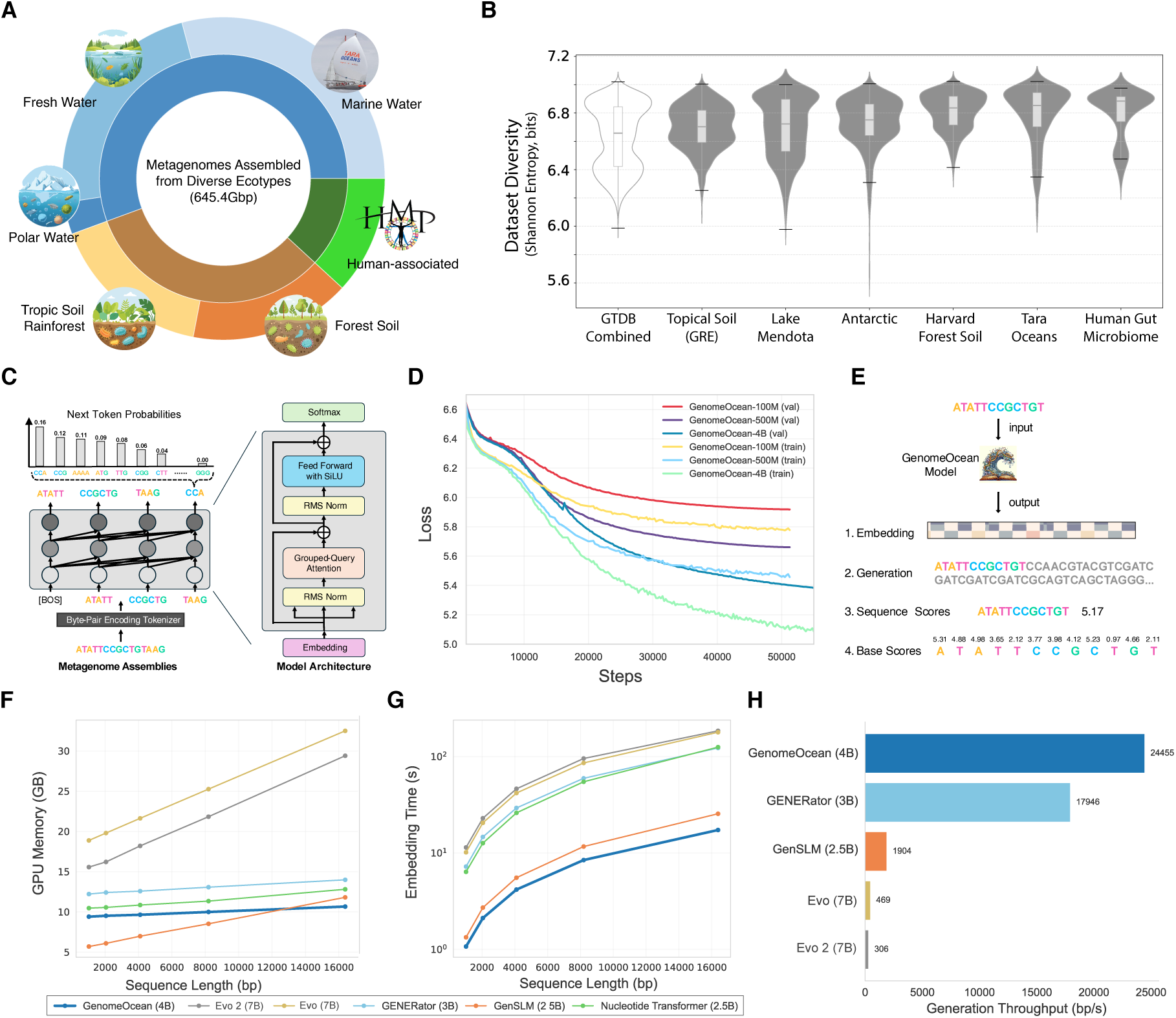
GenomeOcean utilizes global metagenomic diversity and efficient architecture for large-scale genomic modeling. (**A**) Ecological breadth of the pre-training corpus. The model was trained on 645.4 Gbp of high-quality contigs assembled from diverse biomes, including marine and freshwater gradients, terrestrial soils (tropical, temperate forest), polar subglacial lakes, and human-associated microbiomes. (**B**) Sequence information density. Comparison of Shannon entropy (calculated via tetra-nucleotide frequencies, TNF) between the GenomeOcean training set and the Genome Taxonomy Database (GTDB). Environmental assemblies consistently exhibit higher sequence entropy than reference genomes, indicating a richer capture of genomic diversity. (**C**) Model architecture and tokenization. GenomeOcean utilizes a byte-pair encoding (BPE) tokenizer to compress DNA into semantic units, feeding a decoder-only Transformer architecture equipped with Grouped-Query Attention (GQA) and Root Mean Square layer normalization (RMSNorm) for stable, efficient autoregressive modeling. (**D**) Pre-training scaling laws. Training and validation loss curves for 100M, 500M, and 4B parameter models demonstrate that increased scale monotonically improves genomic modeling capacity. The transient spike in the 4B trajectory reflects optimizer re-initialization during training curriculum adjustments. (**E**) Inference modalities. The model supports versatile downstream tasks: encoding sequences into high-dimensional embeddings, autoregressive generation of novel sequences, and per-token or whole-sequence likelihood scoring (perplexity). (**F**–**H**) Computational benchmarks. GenomeOcean demonstrates superior efficiency compared to existing genome foundation models. It achieves lower GPU memory footprint during embedding (**F**), faster embedding latency (**G**), and orders-of-magnitude higher generation throughput (**H**), exceeding 24 Kbp/s.

The resulting corpus creates a dataset integrating freshwater communities from Lake Mendota (Oliver et al., 2024), marine gradients from Tara Oceans (Bork et al., 2015; Sunagawa et al., 2020), human-associated microbiomes (HMP) (Peterson et al., 2009), tropical soil (GRE) (Riley et al., 2023), temperate forest soil (Harvard Forest) (Alteio et al., 2020), and cryophilic communities from Antarctic subglacial lakes (Wang et al., 2019), all co-assembled with the distributed metagenome assembler MetaHipMer (Hofmeyr et al., 2020). In aggregate, these six co-assemblies comprise 645.4 Gbp of high-quality contigs derived from 219 TB of raw sequence reads, representing unique biological information previously unavailable to generative modeling.

The assembly statistics show typical characteristics of short read metagenome assemblies (**Table** 1). While N50 values ranging from 0.71 Kbp to 1.65 Kbp indicate that the bulk of the data consists of sub-gene sized fragments, the sheer scale of the corpus ensures a substantial subset of long contigs. This allows a multi-scale training curriculum combining both short and long context windows to traverse the genomic manifold across multiple resolutions, capturing both local coding rules from the fragmented majority and long-range dependencies such as operon organization and gene clusters. Read mapping rates of 87%–95% confirm that each assembled dataset faithfully represents the underlying community diversity.

**Table 1.** Metagenomic datasets comprising the GenomeOcean training corpus. L50 denotes the number of contigs required to cover 50% of the assembly; N50 denotes the sequence length of the shortest contig at 50% of the total assembly length. *\*Tara Oceans, HMP, and Antarctica datasets were assembled in this study; others are from cited works*.

| Dataset | # Raw reads<br>(billions) | Input size<br>(TB) | Assembly<br>L50 ($10^6$) | Assembly<br>N50 (Kbp) | % Reads<br>mapped | Contigs >500bp<br>(millions) | Assembly<br>size (Gbp) |
| --- | --- | --- | --- | --- | --- | --- | --- |
| **Tara Oceans** | 352.1 | 71.6 | 104.1 | 0.85 | 93 | 363 | 323.1 |
| **Lake Mendota** | 72.1 | 24.3 | 20.9 | 1.17 | 94 | 96 | 105.5 |
| **GRE (Soil)** | 22.7 | 8.0 | 9.1 | 1.65 | 88 | 56 | 75.1 |
| **Harvard Forest** | 8.9 | 3.4 | 16.8 | 1.05 | 95 | 71 | 73.0 |
| **HMP** | 453.1 | 98.0 | 22.4 | 0.71 | 87 | 69 | 54.0 |
| **Antarctica** | 29.2 | 9.7 | 1.8 | 1.59 | 94 | 11 | 14.7 |
| **Total / Range** | **938.1** | **215.0** | **1.8–104.1** | **0.71–1.65** | **87–95** | **666** | **645.4** |

To quantify the complexity of this corpus relative to reference genomes, we calculated the Shannon entropy of tetra-nucleotide frequencies (TNF) as a proxy for information density (**Figure** 1**B**, **Methods** 4.2.2). Because microbial species exhibit distinct, conserved oligonucleotide usage patterns (’genomic signatures’, Pride et al. (2003)), higher TNF entropy reflects a richer superposition of biologically distinct signals rather than random noise. We compared 50 Mbp randomly sampled subsets of our environmental assemblies against the Genome Taxonomy Database (GTDB) representative set (Parks et al., 2022). All environmental datasets exhibited higher entropy than the GTDB. This difference is consistent with the notion that reference databases may represent a more limited genomic diversity than is inherent in the global metagenomic corpus.

### A Genome Foundation Model for Scale and Efficiency

We implemented GenomeOcean with a Transformer decoder architecture and a byte-pair encoding (BPE) tokenizer (**Figure** 1**C**, **Methods** 4.1.1). Unlike character-level models, which treat every nucleotide as a distinct computational unit, BPE compresses DNA sequences into variable-length tokens by their co-occurence frequency. This approach reduces the average sequence length by approximately five-fold compared to character-level tokenization, allowing the model to process longer genomic contexts with reduced computational overhead.

We evaluated scaling behavior by training three model variants (100M, 500M, and 4B parameters) on the above corpus. Consistent with scaling laws in natural language processing (Hoffmann et al., 2022), model performance improved monotonically with parameter count (**Figure** 1**D**). The 4B parameter model exhibited the lowest training and validation losses, indicating a higher capacity to internalize genomic complexity. Consequently, the 4B model was selected for continued training. To comprehend higher-order genomic structures such as operons and biosynthetic gene clusters, we extended the context window from 1,024 to 10,240 tokens (∼51 Kbp) in a secondary training phase. The resulting model, designated **GenomeOcean**, supports three primary inference modes: high-dimensional sequence embedding, prompt-based autoregressive generation, and likelihood scoring at both sequence and base levels (**Figure** 1**E**).

To optimize inference efficiency, GenomeOcean takes advantage of FlashAttention-2 and Grouped-Query Attention (GQA) (**Methods** 4.1.2). We quantified the efficiency gain (**Methods** 4.3.1) by benchmarking GenomeOcean against five genomic foundation models of comparable scale: Evo 2-7B (Brixi et al., 2025), Evo-7B (Nguyen et al., 2024), GENERator-Eukaryote-3B-Base (Wu et al., 2025), GenSLM-2.5B (Zvyagin et al., 2022), and Nucleotide Transformer-2.5B-Multi-Species (endoder-only) (Dalla-Torre et al., 2025). In sequence embedding tasks, GenomeOcean required less than half the GPU memory of Evo 2 to process equivalent sequence lengths (**Figure** 1**F**) while achieving higher speed (**Figure** 1**G**). In sequence generation throughput, GenomeOcean demonstrated a distinct advantage over the GenSLM, Evo, and Evo 2 (**Figure** 1**H**).

For example, while Evo 2 generated approximately 300 base pairs per second (bp/s), GenomeOcean sustained speeds exceeding 24,000 bp/s, an acceleration of approximately 80×.

### GenomeOcean Learns a Coarse-grained Universal Genomic Syntax, Capturing Phylogenetic, Cellular, and Epigenetic Contexts

To determine whether GenomeOcean captures the structural geometry of the genomic manifold from unstructured metagenomic data, we tested its embeddings on two basic tasks: reflecting microbial phylogeny and differentiating cellular genomic contexts.

Standard metagenomic binning relies on tetra-nucleotide frequencies (TNF) and coverage profiles to differentiate species (Kang et al., 2015, 2019). We hypothesized that GenomeOcean, by learning a high-dimensional probabilistic syntax of DNA, would capture phylogenetic signals beyond simple composition. We projected DNA fragment embeddings from a simple 10-species synthetic community, the ZymoBIOMICS Microbial Community Standards, into a two-dimensional space, and then used the Adjusted Rand Index (ARI) to evaluate the performance of HDBScan clustering based on these projections (**Methods** 4.2.3). As shown in **Figure** 2**A**, while TNF provides a baseline separation (ARI = 0.79), GenomeOcean embeddings achieves a slightly better resolution (ARI = 0.86). This performance gain is unlikely due to the high dimensionality of the embedding vectors (3,072) or the effect of BPE tokenization, as embeddings from a randomized GenomeOcean control perform poorly (ARI = 0.31). Including a diverse dataset appears to be a key to learn this phylogenetic signal, as we observed that other genome foundation models trained on diverse prokaryotic genomes (e.g., Evo) performed moderately well (*ARI* = 0.68), whereas models heavily weighted toward human or eukaryotic reference genomes (e.g., HyenaDNA, GENERator) performs poorly to distinguish microbial species (**Figure** S1).

**Figure 2.**
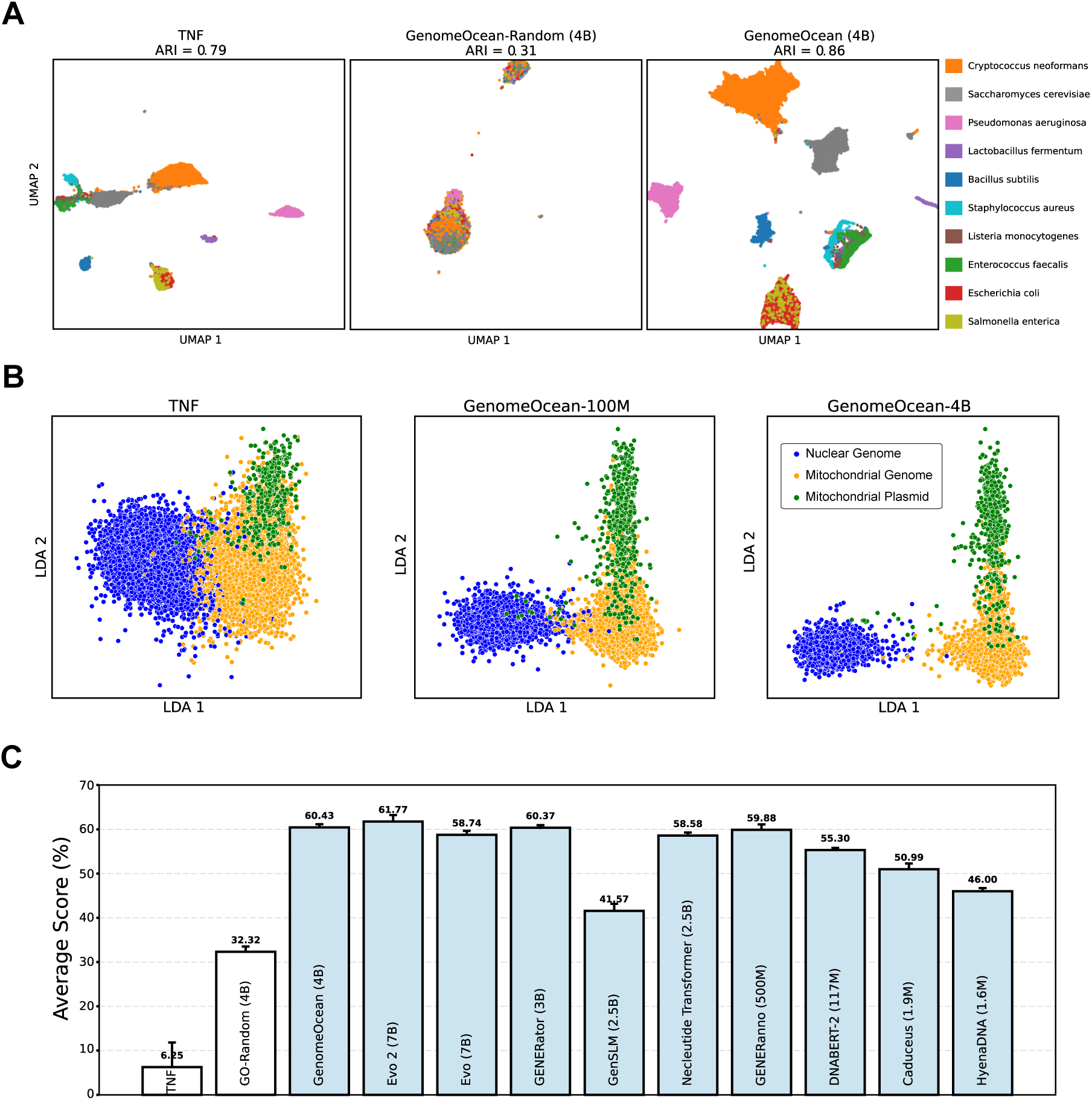
GenomeOcean learns a coarse-grained universal genomic syntax, capturing phylogenetic, cellular, and epigenetic contexts. (**A**) UMAP projections of DNA fragment embeddings from the ZymoBIOMICS Microbial community. GenomeOcean (4B) achieves a higher Adjusted Rand Index (ARI = 0.86) compared to TNF (ARI = 0.79) and a randomly initialized version of GenomeOcean (ARI = 0.31). The model organizes species by their phylogenetic relationship: it forms distinct, proximal clusters for related Firmicutes (e.g., *Enterococcus faecalis*, *Listeria monocytogenes*, and *Staphylococcus aureus*) while merging the highly homologous genomes of *Escherichia coli* and *Salmonella enterica* into a shared space. (**B**) Linear Discriminant Analysis (LDA) projections of fungal sequences categorized as nuclear genomes, mitochondrial genomes, or mitochondrial plasmids. GenomeOcean demonstrates scaling behavior, with the 4B model out-performing the 100M model and the composition-based baseline (TNF). (**C**) Average classification Matthews correlation coefficient (MCC) score on 10 yeast epigenetic-mark prediction tasks. All tested genome foundation models significantly outperform the random baseline (GO-Random, 32.32%), consistent with shared constraints on DNA syntax. GenomeOcean (60.43%) achieves performance comparable to models trained explicitly on eukaryotic reference genomes (e.g., GENERator, 60.37%) and those trained partly on eukaryotic data (e.g., Evo 2, 61.77%).

Consistent with the ARI results, the topology of the projected GenomeOcean embedding space reflects the known phylogenetic relationships. Fragments belonging to distant species form well-separated clusters, whereas those from highly homologous Enterobacteriaceae species *Escherichia coli* and *Salmonella enterica* occupy a shared neighborhood. Fragments from related taxa (e.g., *Enterococcus faecalis*, *Listeria monocytogenes*, and *Staphylococcus aureus*) form nearby but distinct groups. The fungi *Saccharomyces cerevisiae* and *Cryptococcus neoformans*, which belong to different phyla (Ascomycota and Basidiomycota, respectively), also form groups that are distinct both from each other and from bacterial species.

The above experiment suggests that GenomeOcean embeddings improve the “genomic signature” that characterizes microbial species identity, a genome-wide nucleotide composition such as the TNF biases observed in many microbial species (Pride et al., 2003). Another global feature we evaluated is the model’s ability to distinguish sequences based on their cellular context by differentiating nuclear genomes, mitochondrial genomes, and mitochondrial plasmids (**Figure** 2**B**). Identifying mitochondrial plasmids can be challenging due to their genetic diversity, the presence of shared sequences with the main mitochondrial genome, and the complex nature of some mitochondrial genomes (Cahan and Kennell, 2005; Handa, 2008; Kozik et al., 2019). In some cases, the plasmid genome can be integrated into the mitochondrial hosts, creating more challenges for their identification (Smith and Keeling, 2015). We hypothesized that GenomeOcean embeddings would capture subtle, context-specific sequence features distinguishing these replicons, enabling their separation even when their nucleotide composition is similar.

To test our hypothesis, we curated a dataset of 5 Kbp sequences from three genomic classes (**Methods** 4.2.4), and projected their embeddings using linear discriminant analysis (LDA) (**Methods** 4.3.3). The analysis revealed a clear hierarchy of representation capability (**Figure** 2**B**). Tetra-nucleotide frequencies (TNF) failed to distinguish the classes, resulting in overlapping distributions. Support vector machine (SVM) models trained on these embeddings demonstrated that both embeddings from the GenomeOcean 100M and 4B markedly outperformed TNF (**Figure** S2**B**). Specifically, the embeddings achieved a 7-fold reduction in misclassification between mitochondrial plasmids and mitochondrial genomes, while simultaneously improving detection of true mitochondrial plasmid sequences. In addition, GenomeOcean embeddings demonstrated a parameter-dependent emergence of structure: while the 100M model began to separate the classes, the 4B model produced clear, generalizable linear boundaries between sequences derived from these three types of replicons. To validate that this separation arises from learned biological semantics rather than architectural bias, we performed control analyses using LDA and SVM on randomly initialized GenomeOcean baselines (100M and 4B), as presented in **Figure** S2**A** and **C**. These comparisons confirm that the model’s discriminatory capacity results from effective pre-training, as randomized networks fail to recover the clear decision boundaries found in the trained GenomeOcean embedding space.

Finally, we investigated whether the genomic manifold learned by GenomeOcean extends to global regulatory syntax, specifically eukaryotic histone modifications. While our training corpus is predominantly prokaryotic, environmental metagenomes inherently contain genetic material from pico-eukaryotes, fungi, and other microbial eukaryotes.

Eukaryotic gene regulation relies on histone modifications, governed by specific physico-chemical constraints on DNA flexibility and sequence periodicity (e.g., *H3K4me3*, *H3K36me3*) (O’Kane and Hyland, 2019). To assess whether the model can recover eukaryotic regulatory signatures, we evaluated it on ten binary classification tasks predicting yeast epigenetic marks. GenomeOcean (60.43% average Matthews correlation coefficient score) outperformed the random baseline and most foundation models, achieving performance comparable to models trained explicitly on eukaryotic reference genomes (e.g., GENERator) and those trained partly on eukaryotic data (e.g., Evo 2) (**Figure** 2**C**). We show detailed results of 10 tasks in **Tables** S1 and S2.

This suggests that GenomeOcean has learned a universal “regulatory grammar”, likely driven by the fundamental energetics of DNA-protein interactions. While specific chromatin proteins differ between domains, the underlying DNA structural mechanics (e.g., bendability and electrostatic potential required to recruit regulatory complexes) appear to be part of a shared genomic manifold.

### GenomeOcean Learns Fine-Grained Protein-Coding and Evolutionary Constraints

Unlike large eukaryotic genomes, microbial genomes are densely packed with functional coding sequences. However, training a language model solely on raw nucleotides provides no explicit supervision regarding protein structure or function. This challenge is compounded by the use of BPE tokenization, which operates on arbitrary subword units rather than biologically meaningful codons, thereby obscuring the reading frame. Despite these obstacles, we investigated whether GenomeOcean implicitly learns the rules of protein coding and evolution.

We first assessed whether the model could generate a functionally diverse protein repertoire. To evaluate this, we generated a synthetic dataset and annotated the predicted proteins using standard metagenomic tools (**Methods** 4.3.5). Quantifying the number of unique protein superfamilies as a function of sampling depth revealed a discovery curve following Heaps’ law (Heaps, 1978) scaling (**Figure** 3**A**). This unbounded growth indicates that the model has not merely memorized a limited set of common genes but has learned generalizable principles mapping nucleotide patterns to protein structures, enabling the generation of a broad functional diversity of proteins.

**Figure 3.**
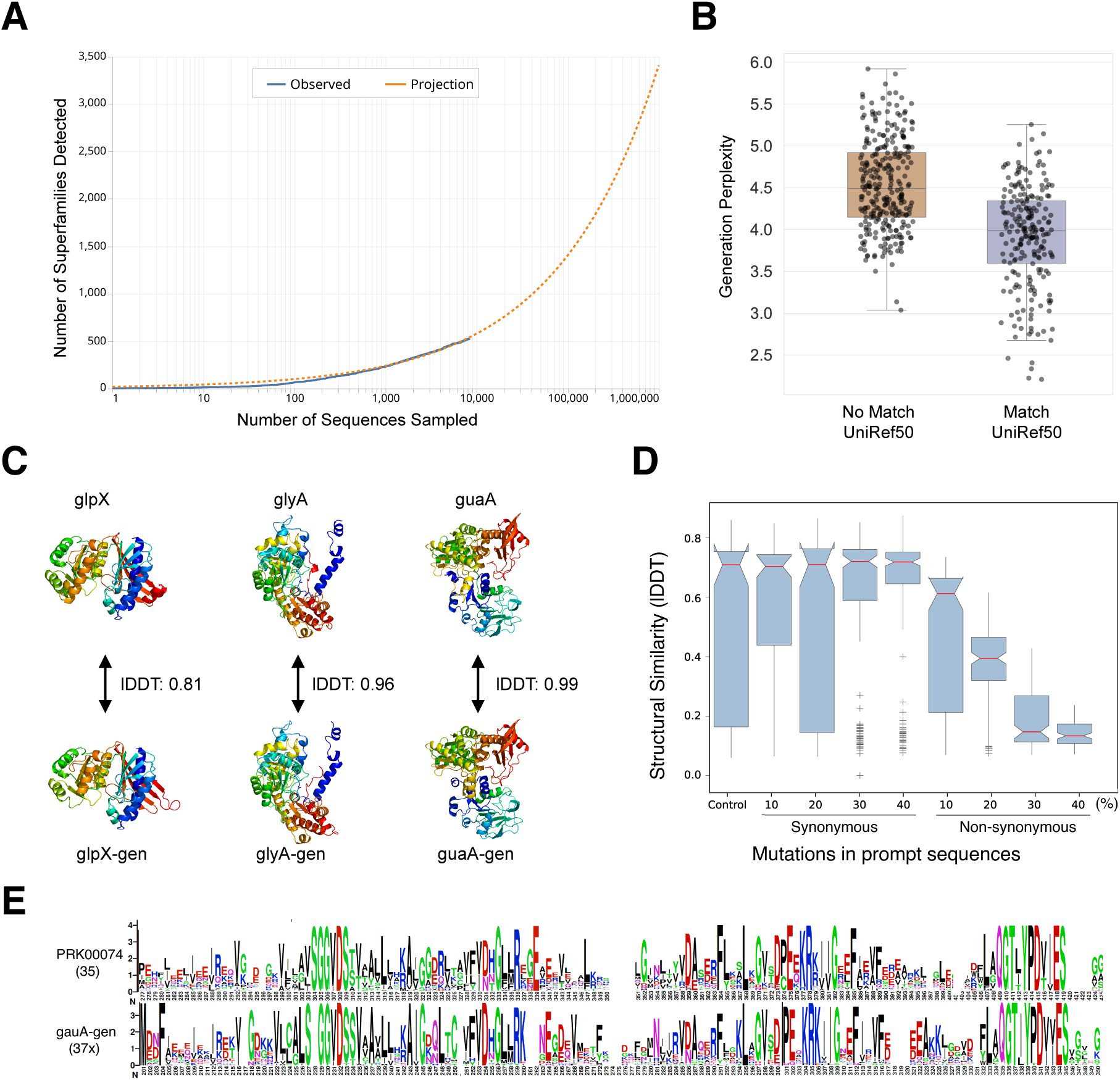
GenomeOcean learns fine-grained principles of protein coding and evolution. (**A**) GenomeOcean-generated sequences recover an expanding diversity of protein superfamilies. The discovery curve and its Heap’s law projection (dashed line) show continued emergence of superfamilies, indicating broad functional coverage. (**B**) Perplexity reflects biological plausibility. During random sequence generation, GenomeOcean assigns consistently higher perplexity scores to “hallucinated” sequences (No Match UniRef50) compared to those with identifiable homologs (Match UniRef50). This separation demonstrates that the model’s internal uncertainty metric can distinguish between biologically supported proteins and hallucinations. (**C**) Gene autocompletion. Using partial *glpX*, *glyA*, and *guaA* sequences as prompts, GenomeOcean successfully completes coding regions. Structural alignment and high local Distance Difference Test (lDDT) scores confirm accurate, functional protein predictions. (**D**) Distinguishing understanding from memorization. To verify that GenomeOcean learns protein semantics rather than memorizing training data, prompt sequences were subjected to synonymous and non-synonymous perturbations. Synonymous codon substitutions result in generated proteins with high structural stability (high lDDT), whereas non-synonymous mutations lead to significant structural divergence. This confirms that the model is robust to silent variation while being sensitive to evolutionarily meaningful changes. (**E**) GenomeOcean captures evolutionary constraints. Sequence logos comparing natural PRK00074 family members with GenomeOcean-generated variants show conserved residue patterns and family-specific signatures, reflecting learned evolutionary information.

Like those trained in other domains, generative genome foundation models also “hallucinated”: generating random sequences without underlying biological functions. We next tested whether GenomeOcean can internally distinguish between biologically plausible proteins and random sequences. We generated 3, 000 sequences with 5 Kbp and extracted the longest open reading frames (ORFs). With the caveat that some ORFs may be entirely novel, we searched these ORFs to identify homology to natural proteins in the UniRef50 database (Suzek et al., 2007) to distinguish plausible versus random generations (**Methods** 4.3.6). We then compared their internal perplexity scores between these two groups. As shown in **Figure** 3**B**, ORFs with valid UniRef50 matches exhibit substantially lower perplexity than those without matches. This separation suggests that GenomeOcean is likely calibrated to biological plausibility: it assigns lower loss to sequences that respect natural protein syntax, effectively discriminating between plausible coding sequences and spurious ones.

How does GenomeOcean recognize protein-coding plausibility? One possibility is that it learns long-range structural dependencies essential for proper folding. To test this, we investigated its ability to generate valid three-dimensional structures from partial proteins. We prompted the model with short gene fragments (300–600 bp) and allowed it to “autocomplete” the remaining coding sequence for three randomly selected structures of varying complexity: *glpX*, *glyA*, and *guaA* (**Methods** 4.3.7). We then folded the generated sequences using Chai-1 (Discovery et al., 2024). Across all three cases, GenomeOcean completed the partial prompts into full-length proteins that recapitulate the native folds, evidenced by high local Distance Difference Test (lDDT) scores (**Figure** 3**C**).

To distinguish whether this performance stems from a deep understanding of protein structure or surface-level memorization of similar proteins in the training data, we probed the model’s sensitivity tsynonymous versus non-synonymous perturbations (**Methods** 4.3.8) in its contexts. Using a sponge-derived TRAP protein as a reference, we mutated the prompt sequence and assessed the structural stability of the generation. The model demonstrated robustness to synonymous codon changes: generating structurally consistent proteins even with 40% synonymous mutation rates, whereas non-synonymous mutations caused immediate structural degradation (**Figure** 3**D**).

As known protein 3D structures were not included in the training data, we postulate that GenomeOcean implicitly learned structural principles by deriving statistics from families with large numbers of diverse members in the metagenomic training data. Consequently, we could sample the learned distribution that reflects evolutionary constraints, to recover such diversity. We compared the sequence diversity of generated *guaA* variants against natural homologs from the PRK00074 family (**Methods** 4.3.9). Sequence logos derived from the generated variants closely match those of the natural family, reproducing conserved residues and specific substitution patterns (**Figure** 3**E**). This agreement demonstrates that GenomeOcean has learned to simulate not just individual proteins, but the evolutionary landscape that constrains them.

Together, these findings show that GenomeOcean, trained exclusively on unannotated metagenome co-assemblies with an unnatural vocabulary, internalizes an implicit grammar of protein coding. It captures fundamental relationships between nucleotide sequences and protein structure, function, and evolution, enabling the zero-shot generation of biologically plausible protein-coding genes. GenomeOcean’s ability to generate diverse yet structurally coherent proteins, separate plausible from implausible ORFs, and reproduce family-level constraints indicates that it is sampling from a structured, low-dimensional coding manifold rather than the full sequence space.

### Cross-Model Convergence on a Shared Genomic Manifold

The **Genomic Manifold Hypothesis** posits that although the space of possible DNA sequences is astronomical (4*^N^*, where *N* is the sequence length), biologically viable genomes exist on a much smaller, lower-dimensional surface (a “manifold”) defined by evolutionary constraints. Genome foundation models may learn the geometry of this manifold by compressing a vast amount of diverse genomic data. In doing so, they organize local features (e.g., motifs, codon patterns) and global features (e.g., composition, regulatory context) into a coherent low-dimensional structure. When a foundation model embeds a sequence, it is placing it along this learned manifold, reflecting its evolutionary constraints (**Figure** 4**A**). If two sequences are close in this space, they share functional potential, even if their sequences are substantially different.

**Figure 4.**
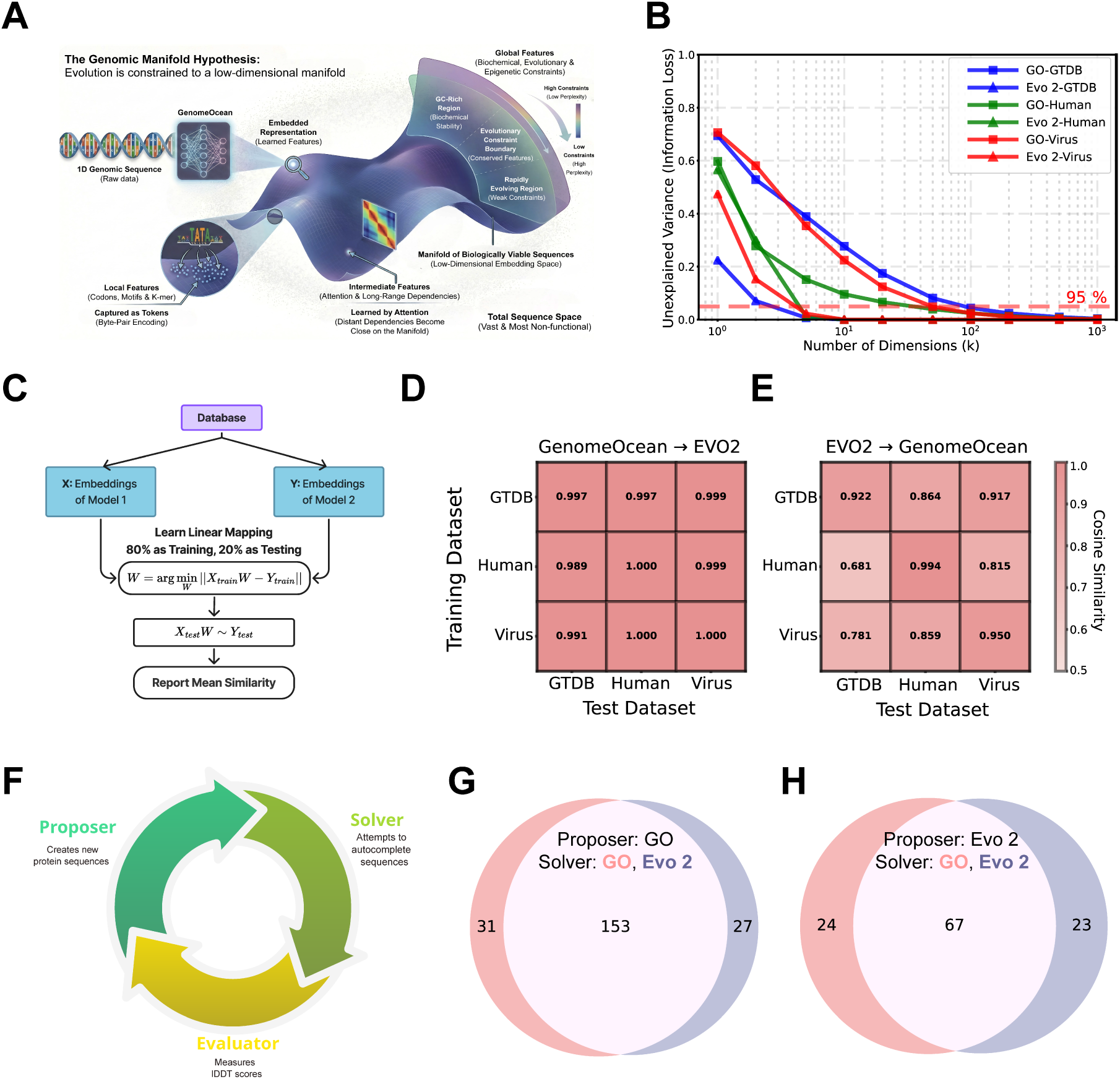
The universal Genomic Manifold. **(A)** Conceptual illustration of the Genomic Manifold hy-pothesis, in which functional genomic sequences occupy a low-dimensional, structured subset of DNA sequence space mapped by GenomeOcean. (Image generated by Gemini 3 using custom prompts and subsequently revised.) **(B)** Analyzing the embeddings of sequences sampled from various datasets shows that most variance in the embeddings can be captured in a small number of principal components, indicating these representations can be compressed onto a low-dimensional manifold. Datasets are designated by different colors: GTDB in blue, Human in green, and Virus in red, while embeddings from GenomeOcean is represented by squares and Evo 2 by triangles. **(C–E)** GenomeOcean and Evo 2 may have learned a common genomic manifold. **(C)** Schematic of learning a linear map *W* between two models’ embeddings using an 80/20 train–test split. **(D–E)** Heatmaps of cosine similarity for GenomeOcean→Evo 2 and Evo 2→GenomeOcean demonstrate that independently trained models learn compatible, linearly alignable representations. **(F–H)** GenomeOcean and Evo 2 may generate protein sequences by sampling a common genomic latent space. **(F)** Proposer–Solver framework: a proposer generates ORFs, a solver shows comprehension by auto-completing them from partial prompts, and an evaluator scores the auto-completed structures using the lDDT metric. **(G–H)** When either GenomeOcean or Evo 2 acts as proposer to randomly generate sequences by sampling their own learned latent space, both models can solve a large shared subset of these sequences, indicating that they may operate within a common region of biologically meaningful coding space and providing convergent evidence for a shared genomic manifold.

If such a genomic manifold exists, it should be uncovered independently by different trained foundation models. We selected GenomeOcean and Evo 2 to test this concept, as their tokenizers (BPE vs single-character tokenizer), model architectures (Transformer vs convolutional multi-hybrid architecture) and training corpus (metagenome assemblies vs reference genomes spanning bacteria, archaea, eukarya, and bacteriophage) are all distinct (Brixi et al., 2025). We designed two complementary analyses to probe their behavior from representational and generative perspectives. For these analyses, we restricted our focus to GenomeOcean and Evo 2, as they were the only models in this study to satisfy two criteria: (i) GenomeOcean and Evo 2 demonstrated superior performance across all embedding-based tasks (**Table** S2). (ii) They were the only models capable of consistently generating sequences that share homology with natural proteins (**Table** S3).

We first asked whether the embeddings of GenomeOcean and Evo 2, despite their high dimensionality (3072 vs 4096), can be compressed into a small number of dimensions, as predicted by the Genomic Manifold Hypothesis. To avoid domain-specific compressions, we examined the effective dimensionality of embeddings from both models across three strategically selected datasets spanning major domains of life: GTDB genomes (bacteria and archaea), Human genomes (eukaryotes), and Viral genomes (**Methods** 4.2.6).

As shown in (**Figure** 4**B**), GenomeOcean embeddings are highly compressible, requiring a relatively small number of principal components to capture most of the variance (e.g., *k* = 87 for 95% variance of the GTDB dataset and *k* = 49 for the virus dataset). The low-complexity human dataset requires even less, *k* = 34. Similarly, Evo 2 retains 95% of its embedding variance with at most *k* = 4 principal components for all three datasets. These results indicate that low-dimensional manifolds exist for the diverse genomic sequences.

We next tested whether the learned manifolds are alignable across models and datasets. Using embeddings from the above GTDB, Human, and Virus datasets, we learned linear maps between the two embeddings (GenomeOcean→Evo 2 and Evo 2→GenomeOcean) by training on 80% of sequences and evaluating the alignment performance on the remaining 20% (**Figure** 4**C**).

As shown in **Figure** 4**D-E**, the linearly projected embedding vectors from one model closely match the embeddings from the other, with high on-diagonal cosine similarity for in-domain projections (e.g., GenomeOcean→Evo 2 on GTDB: 0.997; Evo 2→GenomeOcean on GTDB: 0.922). Cross-domain projections also remain strong (e.g., GenomeOcean→Evo 2 trained on Human, tested on GTDB: 0.989; Evo 2→GenomeOcean trained on Virus, tested on Human: 0.859), supporting a universal latent representation across distinct domains of life.

The surprising observation that Evo 2 compresses the embeddings of rather complex datasets (GTDB, Virus) into only four dimensions brought the possibility of “representation collapse” – a common issue in some pre-trained natural language models where embeddings cluster into a narrow cone in the vector space instead of uniformly distributed (Ethayarajh, 2019). The poorer cross-domain generalization further supports this possibility. To further investigate this possibility, we repeated the above analysis using a randomly initialized GenomeOcean (Random) as a control. Projections from Evo 2→GenomeOcean (Random) yield poor alignment performance and fail to generalize across domains, in contrast to the pre-trained GenomeOcean (**Figure** S3A-B). Although GenomeOcean (Random)→Evo 2 can achieve high in-domain alignment, this is explained by the strong concentration of Evo 2 embeddings around a central vector (**Figure** S3**C**), which allows a linear model to regress any embedding vector to a trivial mean representation upon training, but this linear model does not transfer across domains.

Finally, we investigated whether independently trained models sample from a compatible genomic manifold during generation, indicative of convergent generative behavior. To test this, we employed a Proposer–Solver framework (**Figure** 4**F**), where a proposer model generates 500 random ORFs. The first 300 bp of each ORF serve as the prompt, which the solver model attempts to autocomplete (**Methods** 4.3.12). We defined successful autocompletion as the generation of a sequence whose predicted structure achieves an lDDT score greater than 0.6 relative to the reference ORF (the original ORF proposed by the proposer). We then quantified the intersection of ORFs successfully autocompleted by both GenomeOcean and Evo 2 when acting as solvers on the same set of proposed ORFs. As summarized in **Figure** 4**G–H**, despite significant differences in tokenization, architecture, and training data, the models exhibit substantial cross-model solvability. When GenomeOcean acted as the proposer, both models successfully solved 153 of the same ORFs; conversely, when Evo 2 proposed, they mutually solved 67 ORFs. This overlap indicates that both models operate within a shared, biologically meaningful coding manifold.

## 3 Discussion

The Manifold Hypothesis, a foundational concept in representation learning, posits that high-dimensional natural data (e.g., images, text) concentrate near a low-dimensional non-linear manifold embedded within the ambient space (Fefferman et al., 2016; Narayanan and Mitter, 2010). By effectively mapping the Genomic Manifold, our analysis supports this hypothesis in the context of biology: while the theoretical space of DNA sequences is astronomically large (4*^N^*), evolution has constrained functional sequences to a relatively infinitesimal, structured subset defined by physico-chemical stability and fitness landscapes shaped by evolution. GenomeOcean provides the first scalable, data-driven exploration of this hypothesis. By compressing terabases of raw environmental metagenomic data into a 4-billion-parameter model, GenomeOcean maps a draft topology of this manifold. We demonstrate that the model is able to traverse this latent space by generating valid coding sequences, predicting regulatory syntax, and distinguishing phylogenetic identity. Both GenomeOcean and independently trained models like Evo 2 converge on the ability to “solve” partial ORF prompts (**Figure** 4**C**), supporting that they share a fundamental understanding of local coding constraints.

Models trained on reference databases (e.g., GTDB) are typically constructed to maximize *taxonomic* diversity or based on research interests, often selecting a single representative genome to define a species cluster. While this strategy efficiently spans phylogenetic space, it may inadvertently compress *functional* diversity by filtering out strain-level variation and the dynamic “auxiliary” genes where much of environmental adaptation and specialized metabolism resides. In contrast, training on large-scale metagenomic co-assemblies exposes GenomeOcean to the uncurated “connective tissue” of the genomic manifold, including the vast genetic diversity of the rare biosphere. This deep sampling allows the model to learn the *gradients* of the evolutionary landscape, not just its taxonomic islands. Furthermore, BPE tokenization aggregates nucleotides into semantic units (sub-motifs), allowing the model to allocate its capacity to higher-order syntax (domain organization, gene structure) rather than expending computational resources on spelling out individual nucleotides.

The striking observation that high-dimensional genomic embeddings can be effectively described by a highly compressed low-dimensional manifold suggests that biological sequence space is governed by strict, latent constraints rather than unconstrained stochasticity. Recent findings in protein language models (pLMs), where the intrinsic dimensionality of representations is drastically lower than the ambient embedding space (Valeriani et al., 2022), also support the highly constrained nature of the evolutionary fitness landscape. We propose that these principal dimensions represent the fundamental, independent variables of biological viability—such as physical stability, regulatory logic, and phylogenetic history—distinct from the vast, high-dimensional void of non-functional sequence space. Consequently, this implies that the apparent complexity of genomic data may be reducible to a tractable set of governing rules, offering a geometric framework for both quantifying evolutionary trajectories and engineering novel biological functions within the bounds of viability.

Despite uncovering the low-dimensional geometry of genomic space, our results suggest that GenomeOcean currently captures only the coarse topology of this manifold, not its full resolution. The observed performance degradation in specific abundant protein families (e.g., carbon fixation) and uneven loss across datasets indicates that a 4-billion-parameter model, while efficient, remains under-parameterized relative to the informational density of the global microbiome. While the manifold’s global shape may be low-dimensional, accurately modeling its high-frequency variations requires significantly higher model capacity. With over 25 TB of assembled metagenomes available and petabytes in raw reads from the public sequence repositories, the challenge lies in developing filtering strategies that reduce redundancy without stripping the rare, boundary-defining sequences that give the manifold its structure. Furthermore, reliance on metagenomic assemblies inherently smooths out local manifold texture by collapsing strain-level variation.

Currently, GenomeOcean operates as a “biological autocomplete”, responding to DNA prompts. However, the ultimate utility of the genomic manifold lies in navigation: traversing the latent space to find solutions to specific biological problems. Future work must focus on aligning this generative capability with functional intent, potentially through multi-modal prompting (e.g., conditioning generation on text descriptions, chemical structures, or metabolic goals) or “Best-of-*N* ” sampling strategies constrained by known priors.

In addition to model capacity, several architectural constraints currently limit coverage of the genomic manifold. First, the 50 Kbp context window remains far below the megabase-scale range over which many regulatory and evolutionary interactions operate; modeling full chromosomal or pangenomic scale will require context lengths exceeding millions of base pairs. GenomeOcean’s architecture supports up to 150 Kbp context, but there are only a small number of contigs in the assemblies fall in that size range. Better sequencing technologies, more advanced assembly or scaffolding algorithms are needed to improve assembly continuity. Second, the 4B-parameter model is modest relative to frontier LLMs (Brown et al., 2020; Dubey et al., 2024), indicating that substantially larger or adaptively scaled gFMs will be needed to capture Earth’s full genetic diversity. Third, GenomeOcean uses a dense Transformer rather than a mixture-of-experts (MoE) architecture (Jiang et al., 2024); MoE layers would enable scaling to tens–hundreds of billions of parameters at manageable cost while allowing specialization on rare pathways and niche microbial clades (Dai et al., 2024).

## 4 Methods

### 4.1 Model and Training

#### 4.1.1 Tokenization

Tokenization is the first step of modeling DNA sequences with foundation models. It transforms a DNA sequence into a series of predefined tokens (e.g., CG and AATGC), which are then converted to numerical vectors as input for the foundation model. GenomeOcean adapts a SentencePiece (Kudo and Richardson, 2018) tokenizer with byte-pair encoding (BPE) to tokenize each input DNA sequence into a set of non-overlapping tokens, as proposed in Zhou et al. (2023). The vocabulary of the tokens is determined by iteratively selecting those frequently occurring sequence k-mers (k ranges from 1 to 12 in our case) in the pre-training corpus until the desired size (in this case 4096) is reached. The same tokenizer was used for all the model versions in this work.

#### 4.1.2 Architecture and Pre-training

We optimize the Transformer decoder (Vaswani et al., 2017) architecture with an emphasis on model efficiency. We incorporate FlashAttention-2 (Dao, 2023), which refines the computation process and GPU work partitioning to accelerate attention—the core module of Transformer models. To further improve computational and memory efficiency, we adopt Group Query Attention (GQA) (Ainslie et al., 2023). Unlike standard multi-head attention, GQA partitions query heads into groups, where each group shares a single key head and value head. In addition, we use Root Mean Square (RMS) layer normalization (Zhang and Sennrich, 2019), the Sigmoid Linear Unit (SiLU) activation function (Elfwing et al., 2018) for enhanced representational capacity, and Rotary Positional Embedding (RoPE) (Su et al., 2024) for more flexible positional encoding. During inference, we integrate GenomeOcean with vLLM (Kwon et al., 2023), an efficiency-focused LLM inference library that employs optimizations such as efficient memory management and dynamic sequence batching to increase generation throughput. The model hyperparameters are presented in **Table** 2. GenomeOcean is pre-trained on the Perlmutter supercomputer at the National Energy Research Scientific Computing Center (NERSC)^1^. We scaled the training of the 4B model on 64 NVIDIA A100 GPUs across 16 compute nodes. The first stage costs 14 days, and the second stage costs 1 day. We implemented an efficient multi-node training with Deep-Speed (Rajbhandari et al., 2020).

**Table 2.**
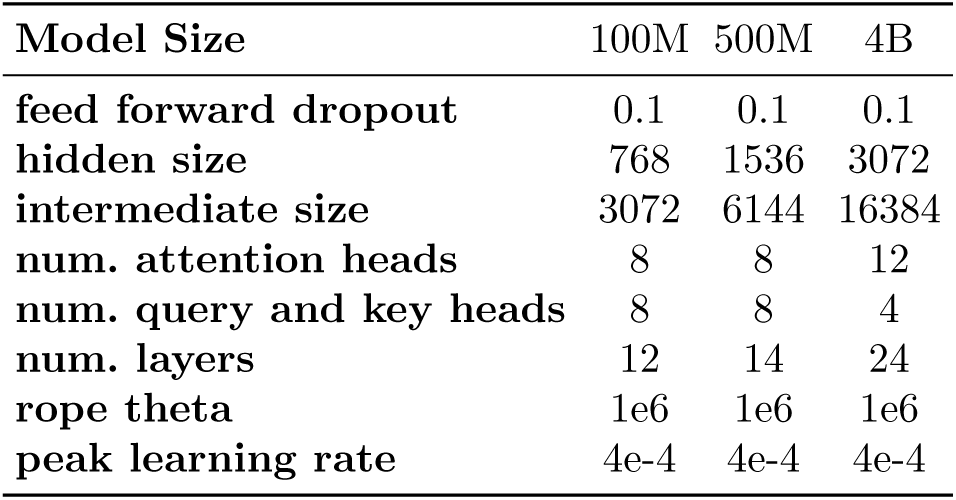
Hyperparameters of GenomeOcean.

| Model Size | 100M | 500M | 4B |
| --- | --- | --- | --- |
| **feed forward dropout** | 0.1 | 0.1 | 0.1 |
| **hidden size** | 768 | 1536 | 3072 |
| **intermediate size** | 3072 | 6144 | 16384 |
| **num. attention heads** | 8 | 8 | 12 |
| **num. query and key heads** | 8 | 8 | 4 |
| **num. layers** | 12 | 14 | 24 |
| **rope theta** | 1e6 | 1e6 | 1e6 |
| **peak learning rate** | 4e-4 | 4e-4 | 4e-4 |

### 4.2 Training and Evaluation Datasets

#### 4.2.1 Assembled Metagenome Datasets

All of the six co-assemblies used for training GenomeOcean were performed using MetaHipMer (Hofmeyr et al., 2020), a distributed metagenome assembler that scales efficiently on supercomputers, enabling it to use hundreds or thousands of compute nodes to co-assemble datasets larger and more complex than what is feasible on shared-memory systems. The assemblies were performed on three national laboratory supercomputers. **Table** 3 provides further details about the assemblies, including resources and time used.

**Table 3.** Assembly details for metagenome datasets used for GenomeOcean training.

| Dataset | Date Assembled | Assembly Time (mins) | Nodes Used | System Used |
| --- | --- | --- | --- | --- |
| **Lake Mendota** | 8/4/2022 | 35 | 1500 | Frontier |
| **Tara Oceans** | 5/10/2023 | 95 | 9000 | Frontier |
| **HMP** | 8/23/2023 | 47 | 9000 | Frontier |
| **GRE** | 11/2/2020 | 84 | 512 | Summit |
| **Harvard Forest** | 11/18/2021 | 11 | 1600 | Cori |
| **Antarctica** | 6/16/2023 | 22 | 1024 | Frontier |

Detailed specifications for each assembly are provided below:

The **Lake Mendota** metagenomes (Oliver et al., 2024) were co-assembled on August 24, 2022, requiring 35 minutes on 1,500 nodes of the ORNL Frontier system. The input consisted of 23.5 TB of FASTQ data containing 153 billion *k*-mers (*k* = 21) with a minimum depth of two. This resulted in 96 million contigs (≥500 bp) totaling 105 Gbp.

The **Tara Oceans** set, comprising 1,211 metagenomes, was co-assembled on May 10, 2023, requiring 95 minutes on 9,000 nodes of the ORNL Frontier system. Inputs consisted of 71.6 TB of FASTQ data across 1,213 samples, containing 1.2 trillion 21-mers with a minimum depth of two. The assembly resulted in 363 million contigs (≥500 bp) totaling 323 Gbp.

The **Human Microbiome Project (HMP)** metagenomes were assembled on August 23, 2023, in 47 minutes on 9,000 nodes of the ORNL Frontier system. The inputs consisted of 98 TB of FASTQ data originating from 13,836 samples, containing 235 billion 21-mers with a minimum depth of two. This resulted in 68.8 million contigs (≥500 bp) totaling 54 Gbp.

The **Great Redox Experiment (GRE)** metagenomes (Riley et al., 2023) were assembled on November 2, 2020, in 84 minutes on 512 nodes of the ORNL Summit system. The inputs comprised 7.7 TB of FASTQ data from 95 samples, containing 86 billion 21-mers with a minimum depth of two. The assembly produced 55.9 million contigs (≥500 bp) totaling 75.1 Gbp.

The **Harvard Forest** metagenomes (Har, 2018; Alteio et al., 2020) were assembled on November 18, 2021, in 11 minutes on 1,600 nodes of the NERSC Cori system. The input used 3.0 TB of FASTQ data from 28 samples, containing 84 billion 21-mers with a minimum depth of two, and resulted in 70.7 million contigs totaling 73.0 Gbp.

The **Antarctic Subglacial Lakes** metagenomes were assembled on June 16, 2023, in 22 minutes on 1,024 nodes of the ORNL Frontier system. Inputs were 9.7 TB of FASTQ data from 21 samples (from FRY, BON, MAT, and WGS), containing 28 billion 21-mers with a minimum depth of two. This resulted in 11 million contigs totaling 14.6 Gbp.

#### 4.2.2 Evaluation of Dataset Complexity

To quantify the diversity in our training corpus, we subsampled each dataset, along with the GTDB (Parks et al., 2022) version R226 representative genomes dataset (including 143,614 species of bacteria and archaea from 189 phyla), to approximately 50 Mbp using BBTools (Bushnell, 2022) version 39.43 [reformat.sh - Xmx110g samplebasestarget=50000000], computed tetranucleotide frequencies (TNF) over 10 Kbp windows using BBTools version 39.43 [tetramerfreq.sh -Xmx110g step=2500 window=10000], and calculated each TNF measurement’s Shannon Entropy using the “entropy” function from scipy.stats version 1.16.0. The violin plots in (**Figure** 1**B** show the distribution of entropies from each dataset, and indicate that each of the large-scale metagenome co-assemblies in our training corpus exceeds GTDB in this measure of diversity.

#### 4.2.3 ZymoBiomics Microbial Community Standard Datasets

The ZymoBiomics Microbial Community Standard (Zymo Research Corp., Irvine, CA, United States), referred to as Zymo dataset, contains the following 10 species: *Pseudomonas aeruginosa*, *Escherichia coli*, *Salmonella eterica*, *Lactobacillus fermentum*, *Enterococcus faecalis*, *Staphylococcus aureus*, *Listeria monocytogenes*, *Bacillus subtilis*, *Saccharomyces cerevisiae*, and *Cryptococcus neoformans*. The original WGS reads can be found with the NCBI Accession number: SRX15657751.

#### 4.2.4 Nuclear and Mitochondrial Genome Sequences Datasets

For the purpose of classification, only published genomes that contain both a mitochondrial and nuclear scaffold were considered. For each published genome, the mitochondrial scaffold and the largest nuclear scaffold were collected for the classification task and classified as mitochondrial genome (MITO) and nuclear (NUCL), respectively.

To collect examples of mitochondrial plasmids, we used genomes from Mycocosm (Grigoriev et al., 2014). In cases where the mitochondrial plasmids were not manually detected, we reassembled them with Flye v2.9.4 for long reads (PacBio), and SPAdes v4.0.0 for short reads (Illumina). In addition, we collected fungal nucleotide sequences from GenBank, whose description field contained different iterations of the string “mitochondrial AND plasmid”. Each genome assembly was then screened for the presence of mitochondrial core hidden Markov model models, such contigs were filtered out. Using Prodigal (Hyatt et al., 2010), potential genes were identified from the remaining contigs, and each potential gene was screened for the presence of DNA/RNA polymerase or reverse transcriptase sequences commonly found in mitochondrial plasmids using a custom hidden Markov model built with HMMer v.3.4^2^. All contigs identified as hits were then saved as mitochondrial plasmid (MITP) contigs.

#### 4.2.5 Epigenetic Mark Prediction Datasets

We used the ten yeast epigenetic-mark prediction datasets^3^ from the Genome Understanding Evaluation benchmark (Zhou et al., 2023). Each dataset corresponds to a binary classification task for one histone modification: *H3*, *H3K14ac*, *H3K36me3*, *H3K4me1*, *H3K4me2*, *H3K4me3*, *H3K79me3*, *H3K9ac*, *H4*, and *H4ac*.

Each sample consists of a 500 bp sequence. All sequences contain only A/T/C/G nucleotides; entries with ambiguous bases were excluded. Following the original protocol, each dataset was randomly split into training, validation, and test sets using an 8 : 1 : 1 ratio. These ten tasks form a standardized, moderately challenging benchmark for evaluating a model’s ability to capture regulatory sequence patterns.

#### 4.2.6 GTDB, Human and Virus Datasets for Cross-Model Convergence Analysis

To evaluate cross-model convergence between GenomeOcean and EVO2, we constructed three benchmarking datasets drawn from phylogenetically and functionally distinct sources.

##### GTDB

We sampled 25,000 genomic segments from the Genome Taxonomy Database (GTDB) representative genome collection (version R226) (Parks et al., 2022), which spans 143,614 bacterial and archaeal species across 189 phyla. Each segment is at most 5 Kbp in length and extracted randomly from the representative assemblies. We used 20,000 sequences for training the linear projection models and 5,000 for testing.

##### Human

To provide a eukaryotic contrast, we extracted 25,000 non-overlapping 5 Kbp segments from the human reference genome (hg38) (Schneider et al., 2017). As with the GTDB dataset, 20,000 sequences were used for training and 5,000 for held-out testing.

##### Virus

To provide a viral contrast, we extracted 25,000 segments of length 5 Kbp from the RefSeq viral database (O’Leary et al., 2016). As with the GTDB dataset, 20,000 sequences were used for training and 5,000 were reserved for held-out testing.

### 4.3 Model Evaluation

#### 4.3.1 Model Efficiency Measurement

We benchmarked sequence embedding and generation efficiency on a single NVIDIA A100 80GB GPU. For embedding, we measured (1) peak GPU memory to encode a single sequence with varying lengths, and (2) throughput as the time to embed 128 sequences of varying lengths. For sequence generation, we measured throughput in base pairs per second. Each model received 1 Kbp prompts and generated 1 Kbp continuations, while the batch size was dynamically adjusted to maximize GPU utilization without exceeding memory limits.

#### 4.3.2 Sequence Composition of Zymo Dataset

The 10 genomes in the Zymo dataset were split into non-overlapping 3 Kbp segments, yielding a total of 24,499 sequences. We evaluated embeddings from tetra-nucleotide frequencies (TNF) and a diverse collection of genome foundation models, including GenomeOcean (4B, pre-trained), GenomeOcean (4B, randomly initialized), Evo 2 (7B) (Brixi et al., 2025), Evo (7B) (Nguyen et al., 2024), GENERator-Eukaryote-Base (3B) (Wu et al., 2025), GenSLM (2.5B) (Zvyagin et al., 2022), Nucleotide Transformer-Multi-Species (2.5B) (Dalla-Torre et al., 2025), GENERanno-Prokaryote-Base (500M) (Li et al., 2025), DNABERT-2 (117M) (Zhou et al., 2023), and Caduceus-ph-131k-Seqlen (1.9M) (Schiff et al., 2024), and HyenaDNA-Medium-160k-Seqlen (1.6M) (Nguyen et al., 2023).

To assess how effectively these embeddings capture sequence composition structure, we applied UMAP to reduce each embedding to two dimensions for visualization, followed by HDBSCAN (Malzer and Baum, 2020) clustering for quantitative evaluation using the Adjusted Rand Index (ARI). Both UMAP and HDBSCAN were run with default settings.

To ensure fair comparison, embeddings for each model were computed using two strategies: (i) the mean of all non-padding token embeddings, and (ii) the final non-padding token embedding for generative models or the [CLS] token embedding for encoder-based models. All experiments were run with random seeds 14, 28, and 42. For each model, we first identified the embedding strategy that achieved the highest ARI across these six configurations. We then report the mean and standard deviation of the ARI in **Table** S2, computed using the optimal embedding strategy under random seeds 14, 28, and 42. For the UMAP visualizations in **Figure** 2**A** and **Figure** S1, we use this optimal configuration, i.e., the one yielding the best ARI across the six settings.

#### 4.3.3 Mitochondrial Plasmid Identification

We first extracted sequence embeddings for classification. Each collected contig was shredded into non-overlapping 5 Kbp segments and paired with the predetermined label of its originating contig. Each segment was then passed through both GenomeOcean-100M and GenomeOcean-4B to obtain sequence embeddings. As a negative control, we additionally computed tetranucleotide frequency (TNF) vectors for every 5 Kbp segment.

Using the resulting embeddings, we evaluated classification performance with support vector machine (SVM) and linear discriminant analysis (LDA). For visualization, 2D LDA projections were generated using the standard scikit-learn LDA implementation, without cross-validation. For SVM classification, we performed stratified 5-fold cross-validation, where stratification was applied across both the class labels and the genome of origin to avoid data leakage (i.e., preventing segments from the same genome from appearing in both training and test splits). The confusion matrix shown in **Figure** S2 aggregates predictions from the test folds across all splits. All SVM models used the default linear kernel in scikit-learn (v1.6).

#### 4.3.4 Epigenetic Mark Prediction

We evaluated embeddings from TNF and a diverse collection of genome foundation models, including GenomeOcean (4B, pre-trained), GenomeOcean (4B, randomly initialized), Evo 2 (7B) (Brixi et al., 2025), Evo (7B) (Nguyen et al., 2024), GENERator-Eukaryote-Base (3B) (Wu et al., 2025), GenSLM (2.5B) (Zvyagin et al., 2022), Nucleotide Transformer-Multi-Species (2.5B) (Dalla-Torre et al., 2025), GENERanno-Prokaryote-Base (500M) (Li et al., 2025), DNABERT-2 (117M) (Zhou et al., 2023), and Caduceus-ph-131k-Seqlen (1.9M) (Schiff et al., 2024), and HyenaDNA-Medium-160k-Seqlen (1.6M) (Nguyen et al., 2023).

To ensure a fair comparison, we computed embeddings using two strategies: (i) the mean of all non-padding token embeddings, and (ii) the final non-padding token embedding for generative models or the [CLS] token embedding for encoder-based models. HyenaDNA and Caduceus were fully fine-tuned due to their small parameter sizes, while all other models were fine-tuned using LoRA applied to the attention layers (key, query, value, and output projection matrices).

Each model was trained with a single-layer linear classification head projecting the embedding dimension to the number of output classes (2 in our setting). We first performed fine-tuning using three learning rates (3×10^−4^, 3×10^−5^, and 3×10^−6^) with a fixed random seed of 42, and selected the best-performing configuration for each model and task. Using the optimal configuration, we then trained each model with three random seeds (14, 28, 42) to report the mean and standard deviation of the performance. We used the Matthews correlation coefficient (MCC) to measure the performance.

#### 4.3.5 Diverse Protein Superfamily Detection

We used GenomeOcean (4B) to generate DNA sequences with uninformed prompts (single nucleotide A, G, C, or T) using default generation hyperparameters (temperature 1.3, top-*k* = −1, top-*p* = 0.7, presence penalty 0.5, frequency penalty 0.5, and repetition penalty 1.0). Prodigal (Hyatt et al., 2010) was used to predict open reading frames (ORFs) which were subsequently annotated using InterProScan (Jones et al.,

2014). Among a total of 9,417 ORFs predicted, 8,340 were assigned to one of 525 superfamilies. We quantified how the number of unique detected superfamilies increases as more generated sequences are included. A Heaps’ law (Heaps, 1978) was used to fit the cumulative superfamily counts across progressively larger subsets of the generated sequences.

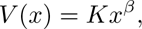

- *V* : The number of distinct superfamiles,
- *x*: The total number of annotated ORFs,
- *K*: The free parameter constant (calibration factor),
- *β*: The exponent parameter determining the growth rate.

In this case, the fitted calibration factor (K) is 16.59, and the fitted rate parameter (*β*) is 0.385407.

#### 4.3.6 Internal Differentiation of Coding Plausibility

To improve the likelihood that GenomeOcean generates sequences matching reference databases, we first performed hyperparameter tuning for sequence generation. We explored *temperature*, *top-p*, *top-k*, and three penalty terms (*presence*, *frequency*, and *repetition* penalties) to identify settings that yield high-confidence outputs. Temperature, top-p, and top-k jointly control sampling diversity. The presence, frequency, and repetition penalties collectively discourage the model from reusing tokens too often, reducing repetitive and low-complexity patterns in the generated sequences.

We applied Bayesian optimization to search this multi-dimensional parameter space. The optimization was guided by two biologically grounded objective functions. The first minimized the proportion of tandem repeats, reducing low-complexity artifacts. The second maximized functional plausibility: generated DNA sequences were translated into proteins, annotated with InterProScan, and scored by the number of proteins containing functional domains covering at least 80% of their length. Both objectives penalized invalid or low-quality generations to steer the search toward robust parameter regimes. This procedure enabled systematic calibration of GenomeOcean’s sampling behavior, yielding generation parameters that enhance the realism and biological utility of synthetic DNA. The optimal set of hyperparameters was:

- temperature: 0.7394,
- top-*p*: 0.5,
- top-*k*: 10,
- presence penalty: 0.9404,
- frequency penalty: 0.1350,
- repetition penalty: 1.5846.

Using the optimized generation hyperparameters, we used GenomeOcean to randomly generate 2,000 DNA sequences, each 5 Kbp long. We identified putative coding regions by running Prodigal (metagenomic mode) to predict nucleotide-level open reading frames (ORFs) and their translated protein sequences. ORFs were then filtered using two criteria: (i) protein length ≥ 100 amino acids; (ii) the amino-acid sequence exceeds a Shannon entropy threshold of 2.5 to remove low-complexity or repetitive sequences.

Next, we clustered all retained ORFs using CD-HIT (Li and Godzik, 2006), which groups protein sequences sharing at least 70% identity using overlapping five–amino-acid words for efficient similarity detection. For each cluster, we selected the longest ORF whose length fell within the desired range (200–400 amino acids, consistent with the upper length limit for reliable structure prediction using ESM-Fold (Lin et al., 2023) on publicly available servers). The resulting collection represents the final set of high-confidence ORFs. From this set, we selected the 500 longest ORFs for downstream analysis.

For each ORF, we computed the perplexity loss and labeled whether it matched a UniRef50 entry, yielding 216 matched and 284 unmatched ORFs. We then compared the perplexity distributions between the matched and unmatched sets, as shown in **Figure** 3**B**.

#### 4.3.7 Gene Autocompletion

We selected three protein-coding genes representing simple, medium, and complex structural complexity. As the simple example, we used the *Shigella flexneri* 2a str. 301 *glpX* gene (Fructose-1,6-bisphosphatase class 2; 1,010 bp). For the medium-complexity example, we used the *Corynebacterium glutamicum* ATCC 13032 *glyA* gene (Serine hydroxymethyltransferase; 1,304 bp). For the complex example, we used the *Staphylococcus aureus* MRSN967703 *guaA* gene (GMP synthetase; 1,541 bp).

For each reference gene, we provided the first 30–40% of the sequence as a prompt: 300 bp, 450 bp, and 600 bp for *glpX*, *glyA*, and *guaA*, respectively. Then we asked GenomeOcean to autocomplete the remaining region. GenomeOcean generated 100 candidate completions per gene, with maximum generation lengths of approximately 1,000 bp, 1,250 bp, and 1,500 bp for *glpX*, *glyA*, and *guaA*, respectively.

The 3D structures of the generated proteins were predicted using ESM-Fold (Lin et al., 2023), and structural similarity to the reference protein was quantified using the local-distance difference test (lDDT). The top-scoring generated sequence for each gene (based on lDDT) was further visualized using Chai-1 (Discovery et al., 2024), as shown in **Figure** 3**C**.

#### 4.3.8 Distinguishing Understanding from Memorization

We evaluated GenomeOcean’s ability to perform gene autocompletion under both synonymous and non-synonymous mutations. For synonymous mutations, a specified percentage of codons was randomly replaced with alternative codons encoding the same amino acid. For non-synonymous mutations, a percentage of codons was randomly altered such that the encoded protein sequence changed.

We used a TRAP transporter solute-binding protein as an example. TRAP is encoded by an uncultured marine bacterium and has a resolved crystal structure (PDB: https://www.rcsb.org/structure/5I5P). A BLAST search using this protein identified a homologous gene from a sponge metagenome (NCBI ID: OY729418), which we used as the reference sequence. The first 500 bp of this gene served as the prompt for mutation analysis.

For each mutation setting, GenomeOcean generated 100 candidate completions, and sequences encoding a valid ORF were retained for evaluation. We used the default generation hyperparameters: temperature 1.3, top-*k* = −1, top-*p* = 0.7, presence penalty 0.5, frequency penalty 0.5, and repetition penalty 1.0. Structural similarity to the reference was quantified using the lDDT score. We tested mutation levels of 10%, 20%, 30%, and 40% for both synonymous and non-synonymous perturbations and visualized the resulting lDDT score distributions in **Figure** 3**D**.

#### 4.3.9 Evolutionary Constraints Analysis of Protein Coding

We used the *Staphylococcus aureus* MRSN967703 *guaA* gene (GMP synthetase; 1,541 bp) as an example to test whether GenomeOcean captures evolutionary-scale patterns. We reused the same 100 autocompleted candidates described in **Section** 4.3.7. Among these, 37 generated sequences encoded a sufficiently long ORF. For comparison, we collected the 35 most diverse natural *guaA* proteins from NCBI^4^. The retained 37 generated ORFs and 35 natural proteins were then used for multiple sequence alignment and protein sequence logo analysis using WebLogo 3 (Crooks et al., 2004).

#### 4.3.10 Effective Embedding Dimensionality Analysis

To assess the effective dimensionality of the embedding space, we conducted a variance-spectrum analysis based on the eigendecomposition of the embedding covariance matrix (Abdi and Williams, 2010; Ethayarajh, 2019). Let *X* ∈ R*^n^*^×^*^d^* denote the embedding matrix, where *n* is the number of sequences and *d* is the dimensionality of each embedding vector. We first mean-centered each embedding dimension by subtracting its average value (*X̄*) across the *N* sequences. We then computed the covariance matrix

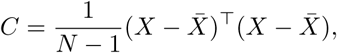

which captures how variance is distributed across the *d* embedding dimensions. We decomposed *C* into its eigenvalues *λ*_1_ ≥ *λ*_2_ ≥ · · · ≥ *λ_d_*, where each eigenvalue reflects the amount of variance explained along a distinct orthogonal direction. This ordered set of eigenvalues constitutes the variance spectrum of the embedding manifold.

To estimate intrinsic dimensionality, we computed the cumulative variance explained by the top *k* compo-nents,

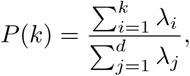

and defined the effective dimensionality as the smallest *k* such that *P* (*k*) ≥ 0.95. We then plotted the unexplained variance (information loss), defined as 1 − *P* (*k*), against the number of dimensions (*k*) on a logarithmic scale (**Figure** 4**B**). This variance-spectrum approach provides an estimate of the manifold’s dimensionality based on how strongly variance concentrates in the leading eigen-directions.

We used three distinct genomic datasets and two models (GenomeOcean and Evo 2) for this analysis:

- **GTDB:** The GTDB testing dataset described in **Section** 4.2.6, consisting of 5,000 sequences.
- **Human:** The human testing dataset described in **Section** 4.2.6, consisting of 5,000 sequences.
- **Virus:** The virus testing dataset described in **Section** 4.2.6, consisting of 5,000 sequences.

We use the mean pooling of all non-padding token embeddings to represent the sequence embedding for both models (GenomeOcean and Evo2) in this analysis.

#### 4.3.11 Geometric Representation Comparison

To assess whether two genome foundation models encode compatible geometric representations, we performed a supervised linear embedding-space mapping analysis.

For the linear mapping analysis, given paired embeddings from a shared dataset, we partitioned the data into 80% training and 20% testing. Let *X* ∈ R*^n^*^×^*^d^*^1^ denote embeddings from Model 1 and *Y* ∈ R*^n^*^×^*^d^*^2^ denote embeddings from Model 2. We learned a linear transformation *W* that best maps one embedding space onto the other by solving

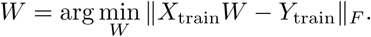

The learned mapping was then applied to the held-out testing embeddings, and cosine similarity between *X*_test_*W* and *Y*_test_ was computed. The mean similarity across the testing set quantifies how linearly recoverable one model’s embedding space is from the other, providing a direct measure of geometric and representational compatibility between the two genome foundation models.

For the linear mapping analysis, we used the same three testing datasets defined in **Section** 4.3.10. The corresponding training datasets used to learn the linear transformation were:

- **GTDB:** The GTDB training dataset described in **Section** 4.2.6, consisting of 20,000 sequences.
- **Human:** The human training dataset described in **Section** 4.2.6, consisting of 20,000 sequences.
- **Virus:** The virus training dataset described in **Section** 4.2.6, consisting of 20,000 sequences.

We use the mean pooling of all non-padding token embeddings to represent the sequence embedding for both models (GenomeOcean and Evo2) in this analysis.

#### 4.3.12 Proposer-Solver Framework Evaluation

We designed a Proposer-Solver framework in which both the proposer and solver can be instantiated by any genome generative model. The proposer first proposes 500 ORFs following the procedure below:

1. Randomly generate 2,000 DNA sequences, each 5 Kbp in length. We identified putative coding regions by running Prodigal (metagenomic mode) to predict nucleotide-level open reading frames (ORFs).
2. Filter ORFs using two criteria: (i) protein length ≥ 100 amino acids, and (ii) amino-acid Shannon entropy ≥ 2.5 to remove low-complexity or repetitive sequences.
3. Cluster the remaining ORFs using CD-HIT (Li and Godzik, 2006), grouping protein sequences with ≥70% identity using overlapping 5 amino acid words. For each cluster, select the longest ORF within the target length range (200–400 amino acids, consistent with the upper length limit for reliable structure prediction using ESM-Fold (Lin et al., 2023) on publicly available servers).
4. Retain the longest 500 ORFs as the proposer’s final set.

For each ORF proposed by the proposer, we extracted the first 300 bp as the prompt and evaluated the solver’s autocompletion ability:

1. To prevent a model from trivially solving its own proposed sequences via memorization, we applied 30% synonymous mutations to the 300 bp prompts.
2. The solver generated 100 candidate completions per prompt. For each, we predicted protein structures using ESM-Fold (Lin et al., 2023) and computed lDDT relative to the reference ORF.
3. The maximum lDDT among the 100 candidates was taken as the solver’s score for that prompt.
4. We counted how many prompts could be autocompleted above the lDDT threshold of 0.6. Alignments in this range (*>* 0.6) strongly correlate with established structural quality metrics and indicate reliably meaningful fold-level agreement (Gilchrist et al., 2024).
5. Independently, we annotated each proposed ORF as matching a UniRef50 or InterProScan entry.
6. We reported the overlap between (i) ORFs that could be autocompleted by the solver and (ii) ORFs that match a UniRef50 or InterProScan entry. This overlap score is the final metric for each proposer-solver configuration.

We evaluated 2 large genome generative models as proposer/solver candidates, including GenomeOcean (4B) and Evo 2 (7B) (Brixi et al., 2025).

##### Generation hyperparameters for the proposer

For GenomeOcean (4B), we used the optimal hy-permeters identified in Section 4.3.6. For Evo 2 (7B), we used the default hyperparameters (temperature 1.0).

##### Generation hyperparameters for the solver

For Evo 2, we reused the same decoding hyperparameters as in the proposer stage. For GenomeOcean, however, we performed systematic hyperparameter tuning. Because tuning Evo 2 would be computationally prohibitive, and the goal of the Proposer-Solver experiment is to explore the shared “common sense” across different models rather than compare their performance, it is reasonable to tune only GenomeOcean. The tuning procedure was:

1. Select the first 100 ORFs proposed by GenomeOcean.
2. Use the first 300 bp of each ORF as the prompt; generate 10 candidate completions per prompt.
3. Perform a grid search over generation hyperparameters: temperature ∈ {0.5, 0.6*, . . . ,* 2.0}, top-*p* ∈ {0.5, 0.6*, . . . ,* 1.0}, and top-*k* ∈ {50, 100, 150*, . . . ,* 400}, holding other hyperparameters fixed.
4. Compute lDDT scores and count ORFs whose maximum lDDT *>* 0.8 (a stringent threshold indicating near-residue-level similarity).

Based on this tuning, we selected the optimal GenomeOcean hyperparameters: temperature 1.2, top-*p* = 0.5, and top-*k* = 100.

## 5 Data and Materials Availability

All data, code, and materials used in this study are available for reproduction and extension of the analysis. No materials transfer agreements (MTAs) or additional restrictions apply. All accession numbers and download links are listed below.

### 5.1 Public Metagenome Assemblies

Three public metagenome assemblies were downloaded from JGI’s IMG/M databases:

1. Lake Mendota: Freshwater microbial communities from Lake Mendota, Crystal Bog Lake, and Trout Bog Lake in Wisconsin, United States (multi-decadal time-series). IMG Submission ID: 288555, doi:10. 46936/10.25585/60001198.
2. Great Redox Experiment (GRE): Lab enrichment of tropical soil microbial communities from Luquillo Experimental Forest, Puerto Rico (95 samples). IMG Submission ID: 255440, doi: 10.46936/10.25585/60000880.
3. Harvard Forest Soil: Forest soil microbial communities from Barre Woods Harvard Forest LTER site, Petersham, Massachusetts, United States (28 samples). IMG Submission ID: 264502, doi: 10.46936/fics.proj.2016.49483/60006003.

### 5.2 Metagenome Raw Reads Datasets Used in Assembly

The raw read datasets are listed below:

1. HMP: A subset of Human Microbiome Project metagenomes (13,836 samples), the data were described in a previous study (Peterson et al., 2009).
2. Tara Oceans: Multi-depth, world-wide sampling of Ocean environments, constituting the largest modern-day worldwide collection of plankton sampled ’end to end’ around the world (1211 samples) (Bork et al., 2015; Sunagawa et al., 2020).
3. Antarctic: Water samples and derived enriched culture from Lake Fryxell and Lake Bonny in Antarctica (21 samples), described in a previous study (Wang et al., 2019).

### 5.3 Code & Models

GenomeOcean is open source and publicly available at https://github.com/jgi-genomeocean/genomeocean.

The models are publicly available at https://huggingface.co/collections/DOEJGI/genomeocean-pilot-models.

### 5.4 Reproducibility Statement

All data processed or generated in this study (including embeddings, generated ORFs, and evaluation results) are available in the main text, the supplementary materials, or via the public repositories above. All data are available in the main text or the supplementary materials.

## 7 Author Contributions

ZW, HL, ZZ, and WW conceived the research idea. ZW supervised the project. YL provided guidance and support to WW. RR, SH, and RE performed metagenome assemblies. WW built the tokenizer. ZZ designed and trained the models. FL developed inference infrastructure. RMK, FC performed the Antarctic experiment and generated data. ZW, WW, ZZ, MA, XS, SK, GL, SGG, MY, HH, ASS, and LS performed model evaluation. ZW, WW, ZZ, RR, SH, RE, and SK wrote the manuscript with input from all co-authors.

## 9 Acknowledgment

The authors thank Alex Copeland for his feedback throughout this project, and Kurt LaButti for his input on the nuclear genome, mitochondrial genome, and mitochondrial plasmid classification tasks.

## 10 Diversity, Equity, Ethics, and Inclusion

This project reflects collaboration across multiple institutions and career stages, with contributions from researchers of diverse backgrounds. All analyses follow transparent, ethical, and community-beneficial data practices. Although no human subjects were involved, we followed inclusive authorship and responsible data practices and remain committed to equitable participation and responsible scientific conduct.

## Supplementary Material

### A Embedding Representation

We summarize the performance of diverse genome foundation models across two embedding-based evaluations in **Tables** S1 and S2: (i) metagenomic binning on the Zymo dataset and (ii) 10 epigenetic mark prediction tasks. Additional binning visualizations for other models are provided in **Figure** S1.

**Table S1.**
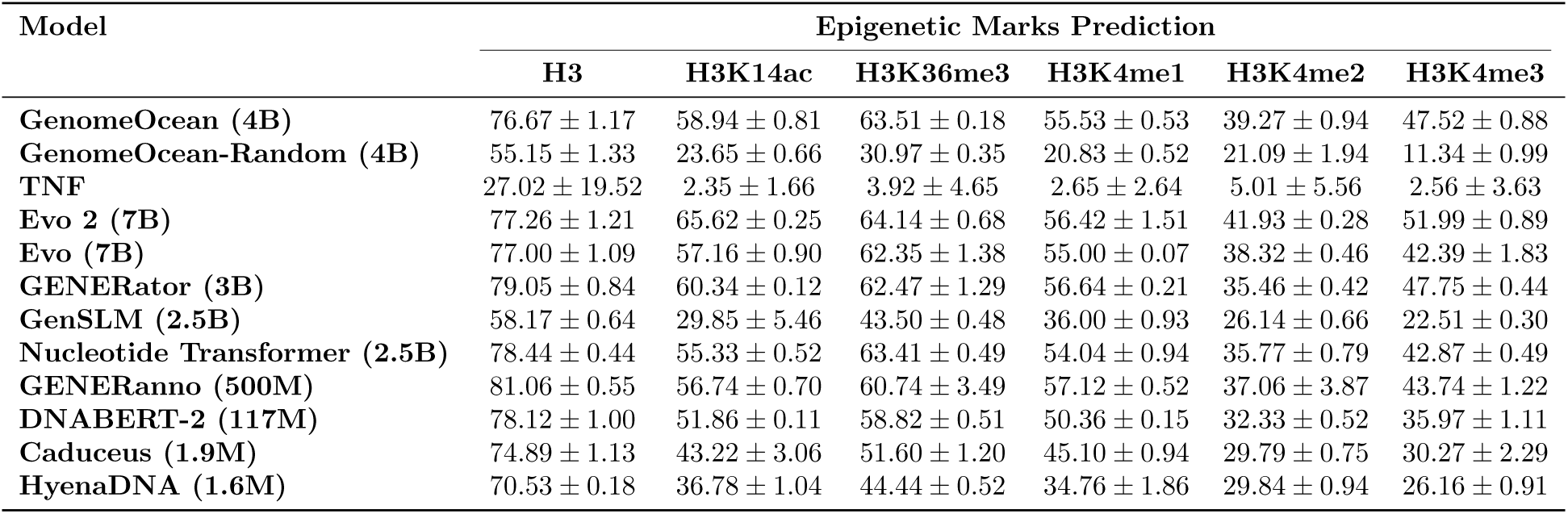
Performance of genome foundation models and TNF across epigenetic mark prediction tasks and metagenomic binning (Part 1). This table reports the classification accuracy (measured by MCC, mean ± standard deviation) of each model on six histone modification prediction tasks.

**Table S2.**
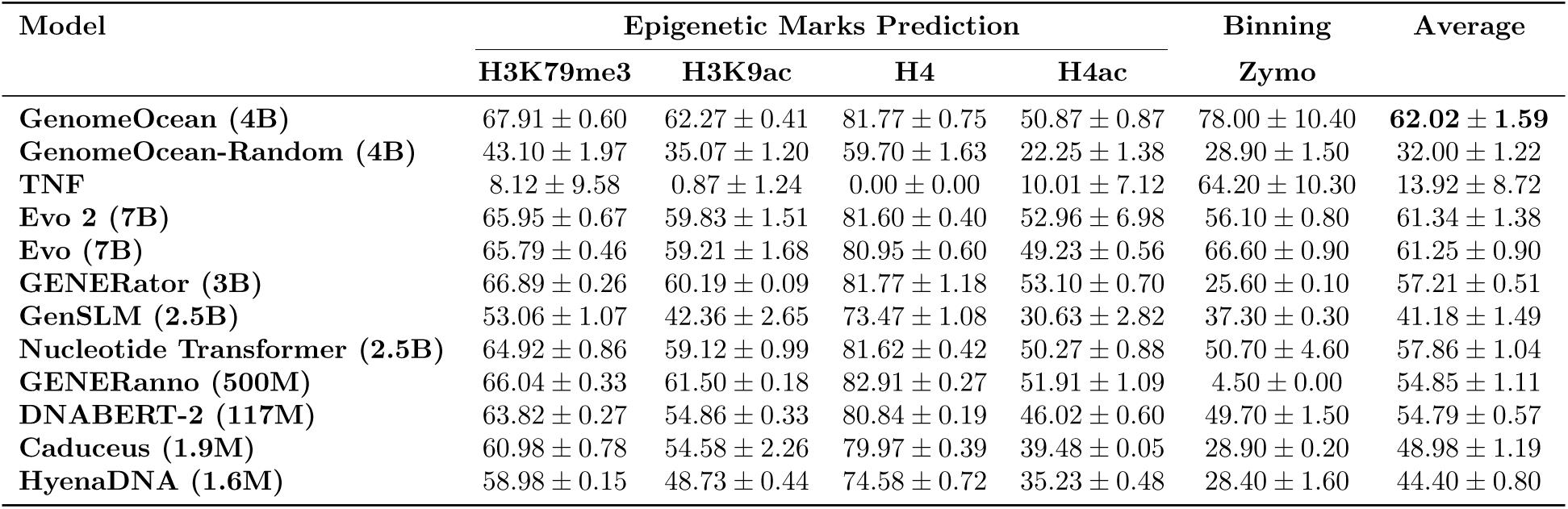
Performance of genome foundation models and TNF across epigenetic mark prediction tasks and metagenomic binning (Part 2). This table presents results on four additional histone marks (measured by MCC) and the Zymo metagenomic binning task (measured by ARI), along with an aggregate average score summarizing overall model performance across all evaluations.

**Figure S1.**
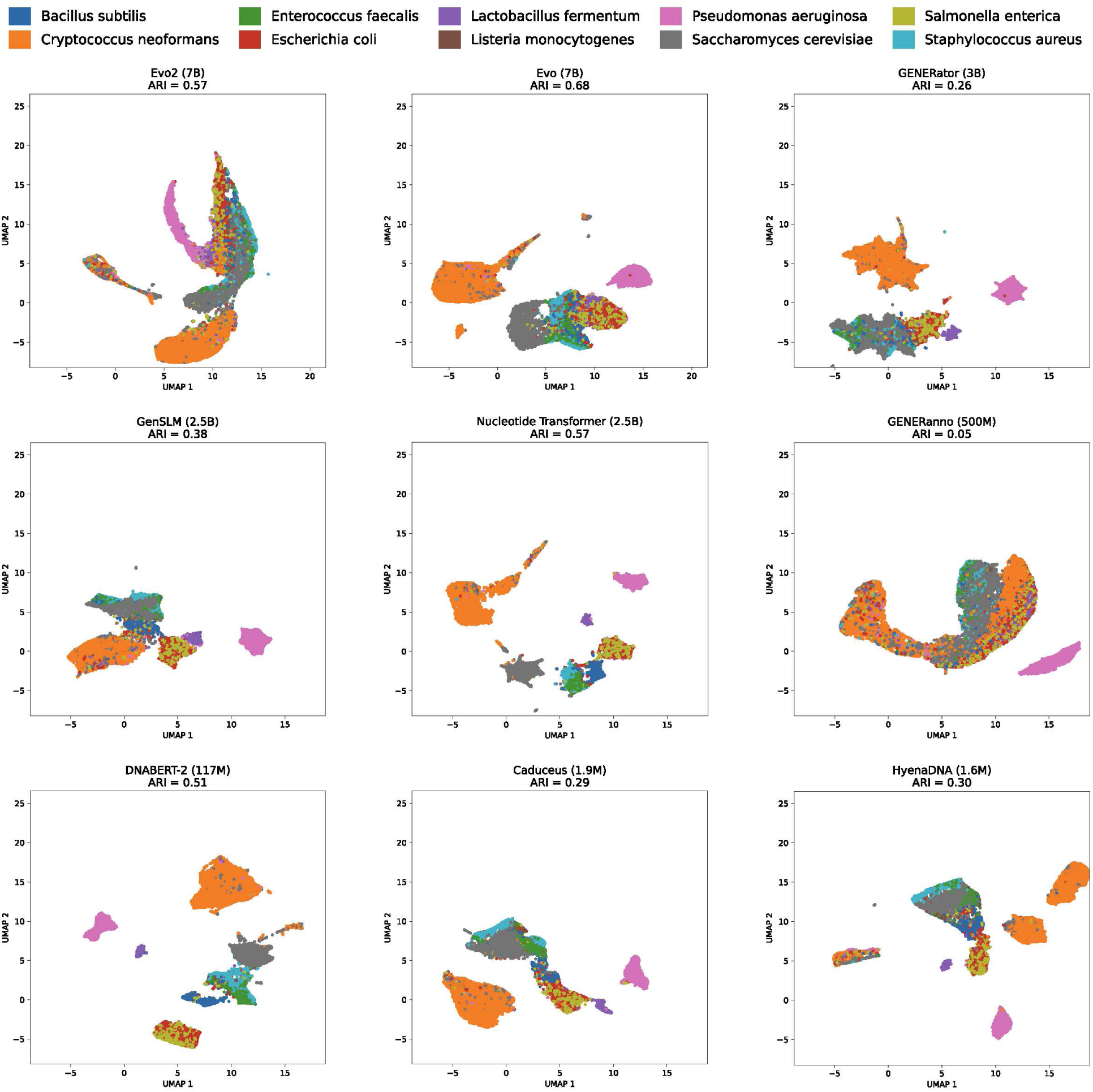
UMAP projections of DNA fragment embeddings from the Zymo dataset using diverse genome foundation models.

### B Control Analysis and Detailed Classification Performance for Genomic Replicons

To validate that the separation of genomic replicons observed in Figure 2B arises from learned biological semantics rather than architectural bias, we performed control analyses using linear discriminant analysis (LDA). In **Figure** S2**A**, we present embedding projections from randomly initialized GenomeOcean baselines (100M and 4B). These comparisons confirm that the model’s capacity to disentangle nuclear (NUCL), mitochondrial (MITO), and plasmid (MITP) sequences results from effective pre-training, as randomized networks fail to recover the clear decision boundaries found in the trained GenomeOcean-4B embedding space. We further quantify this performance using support vector machines (SVM). **Figure** S2**B** details the confusion matrices for the trained models, highlighting the superior separability of GenomeOcean-4B compared to TNF. Corresponding SVM results for the randomly initialized controls are provided in **Figure** S2**C**.

**Figure S2.**
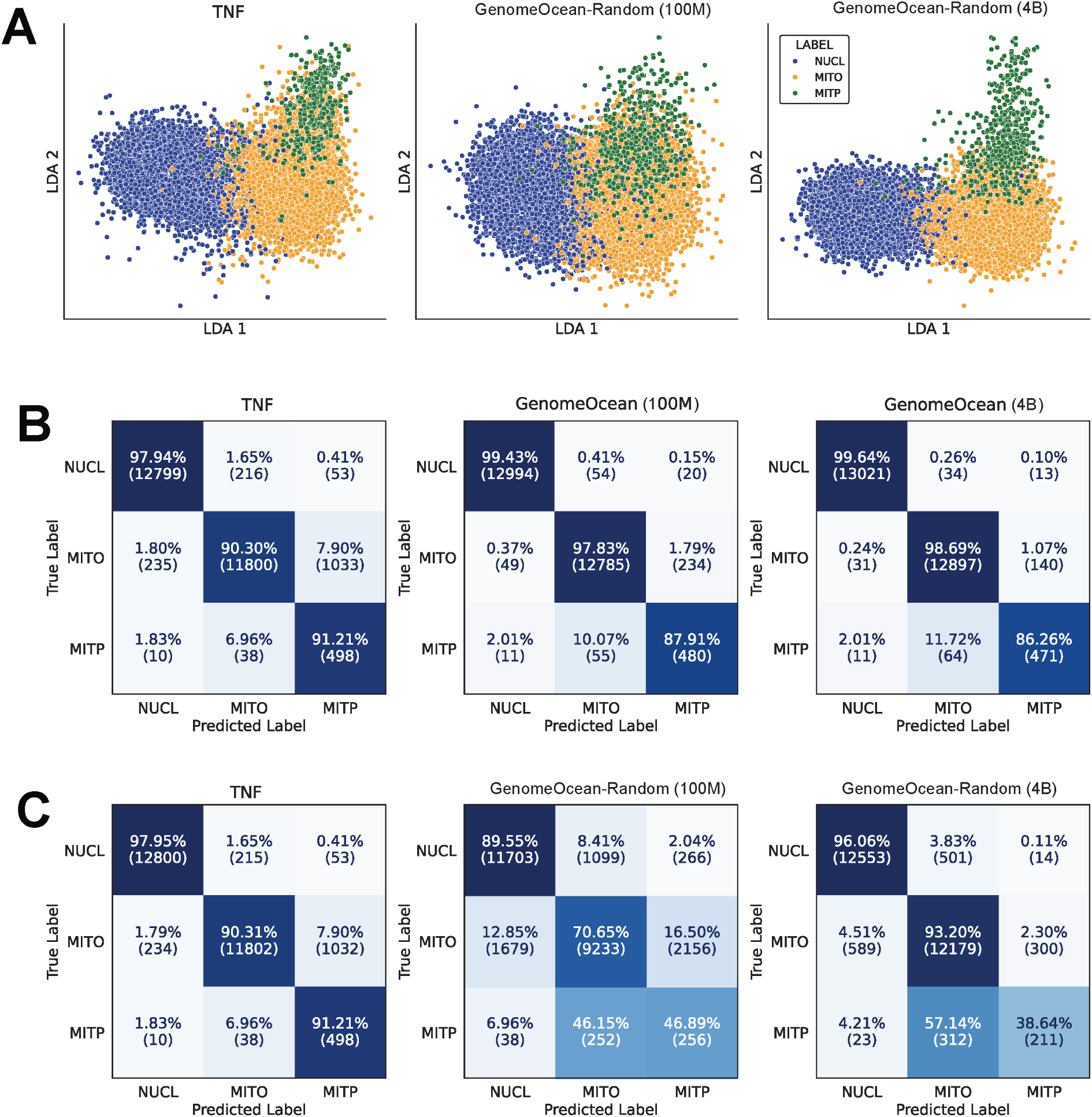
Control analyses and detailed performance metrics for classifying nuclear (NUCL), mitochondrial (MITO), and mitochondrial plasmid (MITP) sequences. (**A**) Linear discriminant analysis (LDA) projections of sequence embeddings from randomly initialized GenomeOcean models (100M and 4B) compared to tetra-nucleotide frequencies (TNF). This serves as a negative control for the results observed in Figure 2B. (**B**) Confusion matrices for classifying nuclear, mitochondrial, and mitochondrial plasmid sequences. Performance of support vector machine (SVM) classifiers trained on TNF, GenomeOcean-100M embeddings, and GenomeOcean-4B embeddings. Rows represent true labels, and columns represent predicted labels. (**C**) Confusion matrices for SVM classifiers trained on embeddings from randomly initialized GenomeOcean models (100M and 4B) compared to TNF, serving as a negative control for the classification performance shown in (B).

### C Model Selection via Natural Protein Homology Analysis

To select suitable models for cross-model convergence analysis, we evaluated the ability of GenomeOcean, Evo 2, Evo, GenSLM, and GENERator to generate sequences homologous to natural proteins. Following the protocol in **Methods** 4.3.6, we generated 2,000 sequences (5 Kbp each) per model, extracted the 500 longest ORFs, and quantified the number of matches against UniRef50 entries.

GenomeOcean utilized the hyperparameters identified in **Methods** 4.3.6, while Evo 2 used default settings. For the remaining models, we performed the following hyperparameter tuning to optimize ORF recovery and minimize sequence repetitiveness, as their default parameters yielded poor performance:

- **Evo (temperature: 1.5):** We evaluated temperatures of 1.0, 1.3, 1.5, and 1.75, ultimately selecting a temperature of 1.5 to mitigate the high repetition observed at default settings. A temperature of 1.5 offered the optimal balance between stability and diversity, yielding 29 ORFs, and 21 of these were sequences with a repetitiveness score (REP) below 40. REP was calculated as the percentage of the total sequence length covered by perfect tandem repeats with a maximum motif size of 7. Lower temperatures (1.3) were ineffective, as they primarily produced highly repetitive sequences rather than diverse biological structures, while higher temperatures (1.75) drastically reduced ORF yield.
- **GenSLM (temperature: 1.3):** We evaluated temperatures of 1.0, 1.3, 1.5, and 1.75, ultimately selecting a temperature of 1.3. The default setting (temperature = 1.0) failed to generate usable ORFs (2 per 100 sequences). A temperature of 1.3 significantly improved recovery (35 ORFs), with negligible marginal gains observed at higher temperatures.
- **GENERator:** We performed a grid search for the generation temperature ranging from 0.5 to 2.0 (step size 0.1). However, the model consistently produced highly repetitive sequences across all tested settings. Consequently, we excluded this model from further evaluation.

The results of the homology analysis are summarized in **Table** S3. For this evaluation, we analyzed the 500 longest ORFs extracted from the 2,000 generated sequences for each model, similar to **Methods** 4.3.6. In instances where a model yielded fewer than 500 total ORFs, all identified ORFs were included in the analysis.

**Table S3.** Number of generated ORFs matching UniRef50 entries. For each model, up to 500 of the longest ORFs were evaluated against the UniRef50 database.

| Model | GenomeOcean | Evo 2 | Evo | GenSLM |
| --- | --- | --- | --- | --- |
| Matching UniRef50 | 216 | 80 | 0 | 0 |

### D Control Analysis of Linear Alignment Stability

For the linear alignment shown in Figure 4C, we include an additional control using a randomly initialized GenomeOcean model to better characterize embedding linear recoverability. GenomeOcean (Random) → Evo 2 (**Figure** S3**A**) exhibits high in-domain similarity but does not transfer across domains, whereas the reverse direction, Evo 2 → GenomeOcean (Random) (**Figure** S3**B**), shows limited transfer both within and across domains.

**Figure S3.**
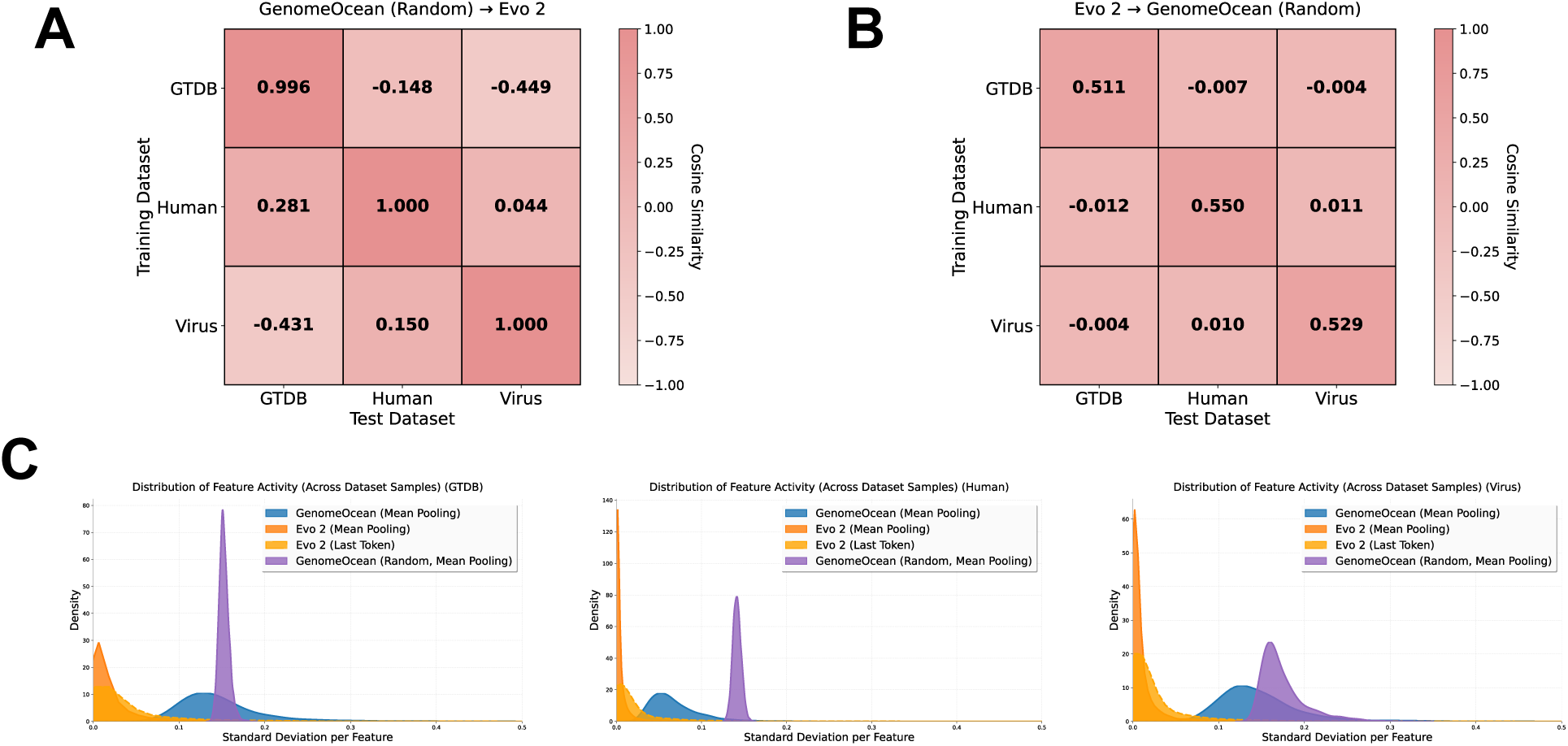
Feature activity and embedding geometry across GenomeOcean and Evo 2. (**A–B**) Linear recoverability controls using a randomly initialized GenomeOcean. Heatmaps show cosine similarity between predicted and true embeddings when learning linear mappings between GenomeOcean (Random) and Evo 2 across three datasets (GTDB, Human, Virus). GenomeOcean (Random) → Evo 2 (A) exhibits deceptively high in-domain similarity but does not transfer across domains, whereas Evo 2 → GenomeOcean (Random) (B) shows poor transfer both within and across domains. (**C**) Kernel density estimates of feature activity for GTDB, Human, and Virus datasets: quantified as the standard deviation of each embedding dimension across samples. Evo 2 (both mean-pooling and last-token embeddings) displays a sharp peak near zero, indicating that most features show minimal variability and concentrate around a common embedding vector. In contrast, GenomeOcean (pre-trained) exhibits a broader distribution, reflecting richer and more informative variation across features. The pronounced low-variance peak in Evo 2 explains why GenomeOcean (Random) → Evo 2 mappings in (A) can achieve high similarity without reflecting true structural alignment.

To understand why GenomeOcean (Random) → Evo 2 yields high in-domain similarity, we quantify feature activity: defined as the per-feature dimension standard deviation of embeddings across GTDB, Human, and Virus datasets (**Figure** S3**C**). Because Evo 2 embeddings exhibit consistent structure across diverse inputs, a linear map can project random embeddings into this space with relatively high similarity, even without reflecting meaningful correspondence. In contrast, Evo 2 → GenomeOcean (Random) does not generalize, indicating that the apparent forward alignment arises from properties of the target embedding space rather than meaningful shared learned features.

## Footnotes

1 https://www.nersc.gov

2 http://hmmer.org/

3 https://www.jaist.ac.jp/~tran/nucleosome/members.htm

4 https://www.ncbi.nlm.nih.gov/Structure/cdd/cddsrv.cgi?uid=234614

